# Axonal spheroids are regulated by Schwann cells after peripheral nerve injury

**DOI:** 10.1101/2024.11.08.622649

**Authors:** Sarah Hunter-Chang, Charlene Kim-Aun, Heeran Karim, Marieke Jones, Tanvika Vegiraju, Ekaterina Stepanova, Brynn Manke, Sarah Kucenas, Christopher Deppmann

## Abstract

Axonal spheroids are hallmark features of neurodegeneration, forming along degenerating axons and contributing to disease progression. Despite their ubiquity across degenerative etiologies, the dynamics of spheroid disappearance, as well as their interactions with glial cells, remain poorly understood. Here, using an *in vivo* zebrafish model of peripheral nerve injury, we identified several patterns of spheroid disappearance that are regulated by Schwann cells. These results describe spheroid dynamics across their lifetimes, establish a role for the extra-axonal environment in altering spheroid outcomes, and identify a cellular mechanism whereby spheroid fates are altered.

**Summary:**

- Axonal spheroids (also called “dystrophic neurites,” “axonal swellings,” and “axonal beading”) undergo three fates: shrinking, breakdown, and uniform disappearance
- Schwann cells ensheath most axonal spheroids and internalize a subset of them
- Schwann cells regulate axonal spheroid fates, identifying a role for axon-extrinsic processes in spheroid regulation

Graphical Abstract:
Following injury, spheroids are more likely to undergo shrinking and breakdown when Schwann cells (SCs) are present on degenerating axons. When Schwann cells are absent, spheroids are more likely to undergo uniform disappearance. Spheroids expose phosphatidylserine, which could facilitate interactions between spheroids and SCs.

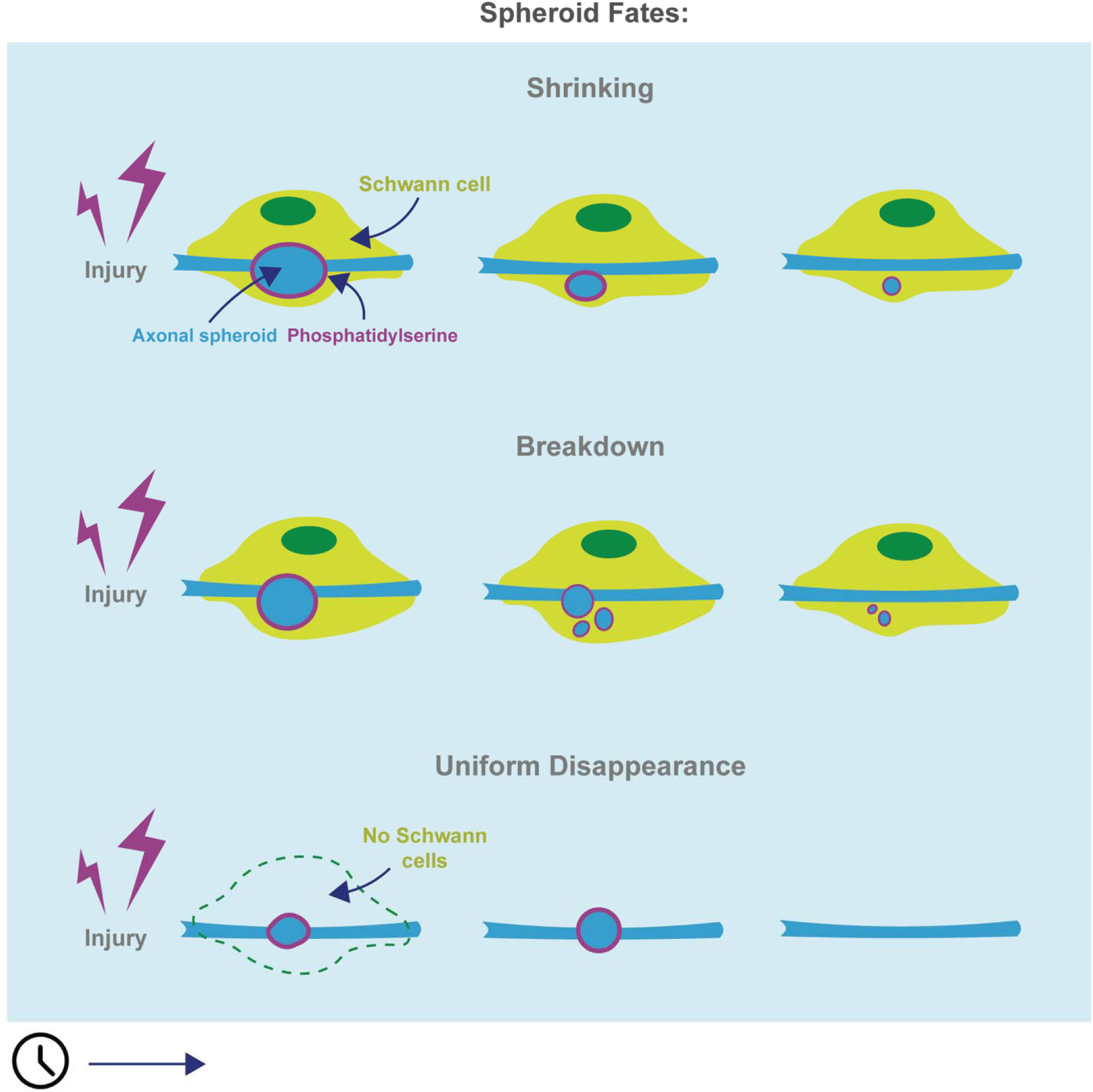

## Introduction

Axon degeneration is a key process in normal developmental circuit refinement, but it also characterizes and drives neurodegenerative disease <u>[1–5]</u>. While any number of different triggers such as physical injury, metabolic stress, or genetic mutations can induce axon degeneration, all etiologies ultimately converge on one of two programmed death pathways <u>[6]</u>. The first pathway is mediated by caspases activated in response to trophic factor deprivation and apoptotic signals, while the second involves the NADase SARM1, which is triggered by axonal NAD+ depletion due to transport disruption or metabolic disturbances <u>[7–10]</u>. Although the specific molecular pathways activated vary by context, both pathways share common cellular features during axonal death <u>[6]</u>. Understanding these shared features is critical for advancing our knowledge of both axon-specific degeneration and broader neurodegenerative processes.

Axon degeneration progresses through two stages: latency, a quiescent stage after the degenerative trigger in which axons remain structurally and functionally intact, and catastrophic degeneration, one of rapid, irreversible disintegration <u>[2]</u>. The formation of axonal spheroids, bubble-like protrusions also known as axonal swellings, beading, or dystrophic neurites, is a hallmark event during the transition from latency to irreversible degeneration <u>[11]</u>. Since the initial description of spheroids over a century ago by Santiago Ramón y Cajal <u>[12]</u>, numerous studies have explored spheroid formation and composition. Spheroids are thought to form due to calcium-driven osmotic water influx and cytoskeletal rearrangement <u>[13–15]</u>, and they often contain pathology-associated substances such as protein aggregates, cytoskeletal elements, aberrant organelles, and excess calcium <u>[11,13,14,16–18]</u>. More recently, studies demonstrate that spheroids can impede action potential conduction as they grow <u>[19]</u> and may rupture to release pro-degenerative factors, potentially exacerbating neural damage <u>[15]</u>.

Despite advances in understanding spheroid formation and function, spheroid fates at later stages of degeneration remain largely unknown. Most prior studies have utilized *in vitro* and *ex vivo* models, limiting insights into spheroid interactions with the native environment. This is significant because axons and glial cells are structurally and functionally intertwined *in vivo*. In the peripheral nervous system, Schwann cells (SCs) ensheath axons and play essential roles such as calcium regulation and metabolic support <u>[20–22]</u>. Given that spheroid formation relies on factors like calcium levels and NAD+ loss, SCs may alter spheroid dynamics <u>[23]</u>. Moreover, SCs robustly respond to axonal degeneration by clearing nerve debris via phagocytic receptors that recognize the ‘eat me’ signal phosphatidylserine (PS) <u>[24,25]</u>. Intriguingly, we and others have observed that spheroids expose PS on their outer membranes *in vitro* <u>[26,27]</u>. These observations suggest that spheroids may interact with SCs in the extra-axonal environment, potentially altering spheroid fate. Therefore, we aimed to characterize spheroid dynamics *in vivo* and determine how interactions with the extra-axonal environment, particularly with SCs, influence their fate.

Here, using zebrafish as an *in vivo* model of axonal degeneration, we describe spheroid dynamics after injury and examine how SCs contribute to spheroid fates. We find that spheroids can persist beyond axon fragmentation and disappear via several mechanisms. Using a complementary *in vitro* model, we determine that these distinct spheroid fates are influenced by the extra-axonal environment. We confirm this *in vivo*, where we find SCs contact most spheroids and are a critical factor mediating spheroid dynamics. Collectively, our results establish the influential role of the extra-axonal environment—specifically SCs—in altering spheroid fate and identify a cellular mechanism underlying spheroid disappearance.

## Results

### Axonal spheroids accumulate during and persist after peripheral nerve injury

Axonal spheroids are calcium-rich focal swellings that form along degenerating axons, appearing as a characteristic "beads-on-a-string" pattern <u>[12,13,26]</u>. Despite advancements in our knowledge of spheroids, we lack understanding of spheroid dynamics beyond their initial formation and of how the extra-axonal environment affects their fates. We therefore established an *in vivo* zebrafish peripheral nerve injury model to study spheroid dynamics in their native environment (Fig. 1a). We chose to laser-axotomize the posterior lateral line nerve (pLLN) in 3-day-post-fertilization (dpf) *NBT:DsRed;HuC:GCaMP6s* zebrafish larvae, allowing unobstructed visualization of nerve degeneration when all glial populations are present <u>[28–30]</u>.

**Figure 1:**
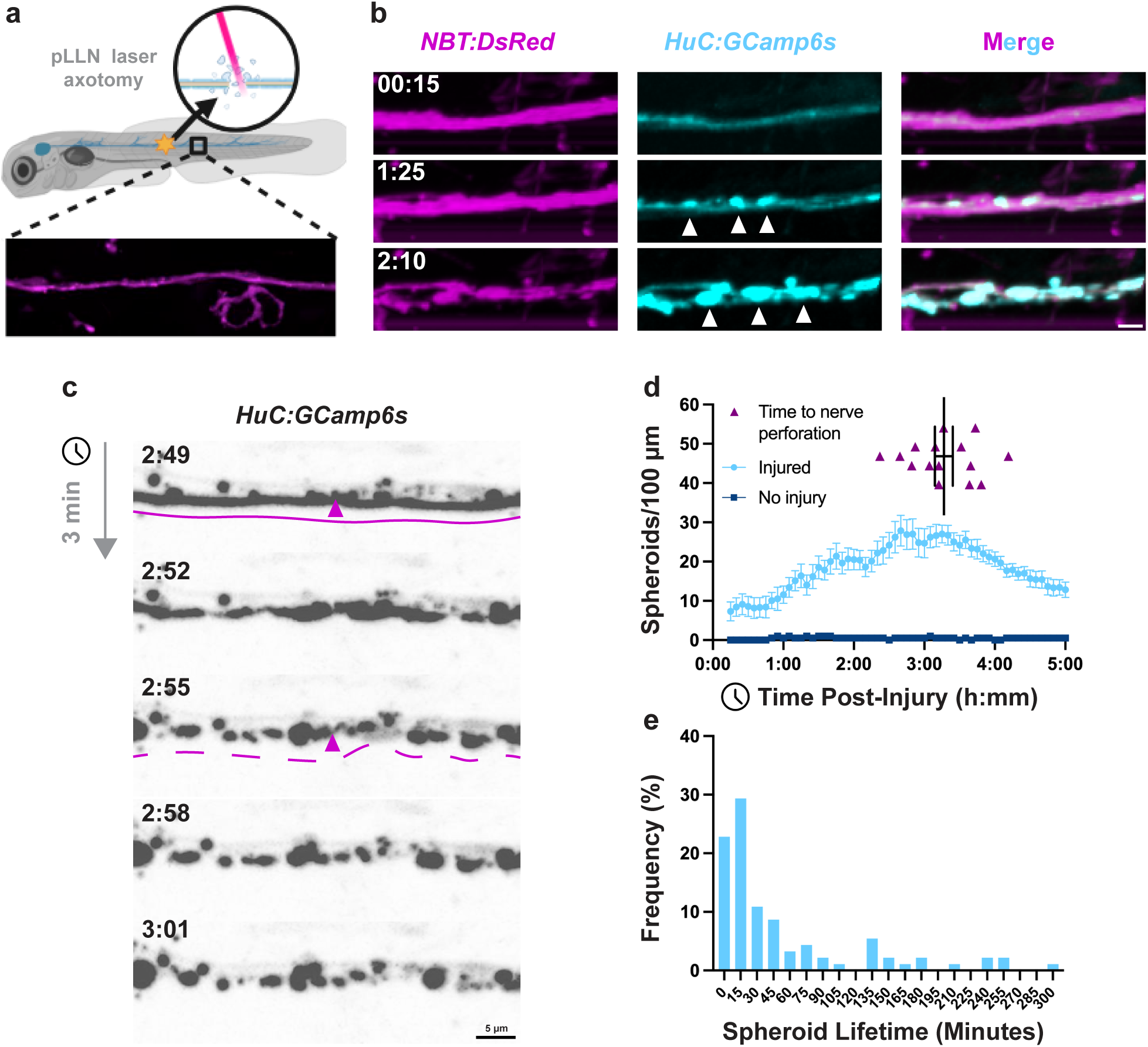
Axonal spheroids are present before and after total nerve perforation. a) A model to study spheroids in vivo: the posterior lateral line nerve (pLLN) of 3dpf zebrafish is transected using laser axotomy, then imaged with time-lapse, confocal microscopy centered 200 μm distal to injury site. b) Axons (*NBT:DsRed*) give rise to calcium-enriched axonal spheroids (*HuC:GCamp6s*, white arrowheads, N=6 fish). c) Calcium transients appearing in pLLN axons after injury (h:mm), followed by axon perforation (solid vs. dashed lines). d) Spheroids as a percentage of maximum quantity over time after injury (N=10 injuries, 2 no injury control fish, error bars = SEM) and times at which the last visible axon is perforated, total nerve perforation (N=15 fish, error bars = SEM). e) Histogram of spheroid lifetimes (minutes, N=92 spheroids from 7 fish).

We transected the pLLN with a pulsed nitrogen-dye laser and used *in vivo*, time-lapse imaging to visualize axonal spheroids in the region 150 to 250 µm distal to the injury site (Fig. 1a). The *NBT:DsRed* transgene labeled the entire pLLN (magenta) as a stable reference for axon morphology (Fig. 1b), while the *HuC:GCaMP6s* transgene allowed visualization of calcium transients and spheroids (cyan) (Fig. 1b). Following pLLN transection, we observed calcium-positive axonal spheroids forming a "beads-on-a-string" pattern along injured axons, with all GCaMP6s signals co-localizing with DsRed (Fig. 1b).

To characterize spheroid dynamics during the later stages of degeneration, we focused on axon fragmentation—the final phase of axon degeneration. In 3 dpf *HuC:GCaMP6s* larvae with injured pLLNs, we observed axon fragmentation (Fig. 1c) and determined the time to total nerve perforation—the point at which all axons in the nerve have fragmented— to be 197 ± 7.5 minutes (mean ± SEM) (Fig. 1d). We validated that the *HuC:GCaMP6s* transgene faithfully represented axon fragmentation by comparing times to total nerve perforation measured with *HuC:GCaMP6s* to those measured with the *NBT:DsRed,* and observed no significant difference (Supplementary Fig. 1).

Using nerve perforation to define completion of axon degeneration, we measured spheroid quantities during and after axon degeneration with a machine learning algorithm in Imaris (Supplementary Fig. 2, Supplementary Materials 1). We counted spheroids from 15 to 300 minutes post-injury and found that spheroid quantities peaked at 160 minutes post-injury and remained elevated until 215 minutes. Subsequently, the spheroid quantities steadily decreased until 300 minutes post-injury (Fig. 1d). Notably, total nerve perforation occurred during the peak of spheroid presence (197 ± 7.5 minutes), indicating that spheroids coincide with ongoing axon degeneration. Furthermore, the continued elevation of spheroid quantities after total nerve perforation demonstrates that spheroids persist even after their axons of origin have fragmented.

To understand individual spheroid dynamics over time, we measured the time from spheroid formation to signal loss, which we refer to as the “lifetime”. We found that spheroids had a median lifetime of 20 minutes, with the plurality (29.3%) persisting between 15 and 30 minutes. Only 28% of spheroids persisted longer than 60 minutes, and 5.43% endured beyond 240 minutes (Figure 1e). Taken together, these studies demonstrate that spheroids are present before, during, and after axon degeneration of the pLLN, but individual spheroids have variable lifetimes that rarely persist for the entire nerve fragmentation process.

### Shrinkage, breakdown, or uniform disappearance define spheroid fates

While quantifying spheroid lifetimes, we observed *HuC:GCaMP6s^+^*spheroids exhibiting several morphological patterns of disappearance. We observed spheroids shrinking (Fig. 2a, Supplementary video 1) and seemingly breaking down into smaller puncta— referred to here as “breakdown” (Fig. 2b, Supplementary video 2). We also observed spheroids disappearing uniformly without apparent changes in morphology—referred to here as “uniform disappearance” (Fig. 2c, Supplementary video 3)—similar to that observed previously *in vitro <u>[15]</u>*. These patterns were also visible using *NBT:DsRed* labeling as well, indicating that spheroid cytoplasm, not just calcium, undergoes these fates (Supplementary Fig. 3).

**Figure 2:**
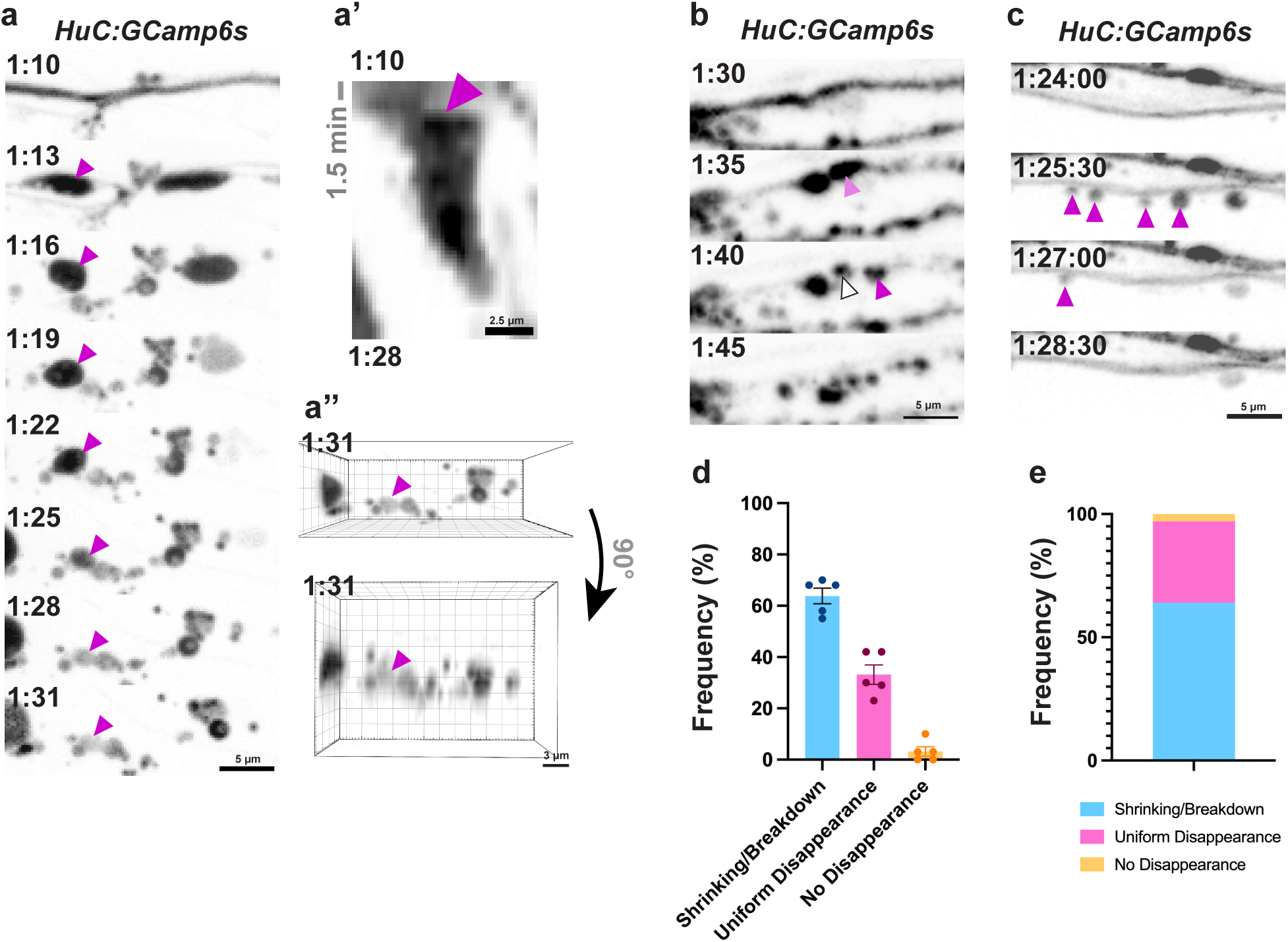
Shrinkage, breakdown, or uniform disappearance defines spheroid fates. a) GCaMP6s-labeled spheroid shrinking (arrowhead) a’) Kymograph showing shrinking spheroid indicated in a. a’’) The ROI in a and an orthogonal view of the ROI in a showing the shrinking spheroid (magenta arrowhead) remains within the z planes captured. b) GCaMP6s-labeled spheroids break down into smaller puncta (arrowheads). c) GCaMP6s-labeled spheroids undergo uniform disappearance, or signal loss with no apparent morphological changes (arrowheads). d) Frequency of spheroid fates plotted by larva and e) averaged together (N=148 spheroids from 5 fish).

To quantify the frequencies of each disappearance pattern, we combined shrinking and breakdown into a single category because they were difficult to distinguish and often occurred together. We found that 64 ± 3.0% and 33 ± 3.8% of spheroids underwent shrinking or breakdown and uniform disappearance, respectively, while 3.2 ± 1.8% did not disappear up to 5 hours post-injury (Fig. 2d, e). We did not observe any obvious relationships between spheroid fates and their location along the pLLn, spheroid size, or signal intensity. These findings identify three distinct morphological fates of spheroid disappearance after formation.

### The extra-axonal environment is required for spheroid shrinking and breakdown

Our previous *in vitro* work investigating spheroids reported only spheroid fates consistent with uniform disappearance <u>[15,26]</u>, whereas *in vivo* we observed the additional fates of shrinking and breakdown described above. We hypothesized these differences were due to the extra-axonal environment, which could give rise to the observed fates via mechanisms such as calcium regulation <u>[21]</u>, phagocytic interactions <u>[31,32]</u>, or breakdown into apoptotic bodies <u>[33]</u>. We therefore tested whether the extra-axonal environment influenced spheroid fates. We harvested superior cervical ganglia (SCG) cells from postnatal day 0*–*2 C57Bl6 mice and plated them in microfluidic devices (MFDs). After 5*–*7 days *in vitro*, neuronal cell bodies were aspirated from their chamber, leaving behind severed axons in the adjacent chamber (Fig. 3a). Live time-lapse confocal microscopy was performed using Fluo-4 AM calcium dye to label axons and spheroids, and cascade blue-conjugated 3kDa dextran to negatively label axons and spheroids <u>[13,26]</u> (Fig. 3b).

**Figure 3:**
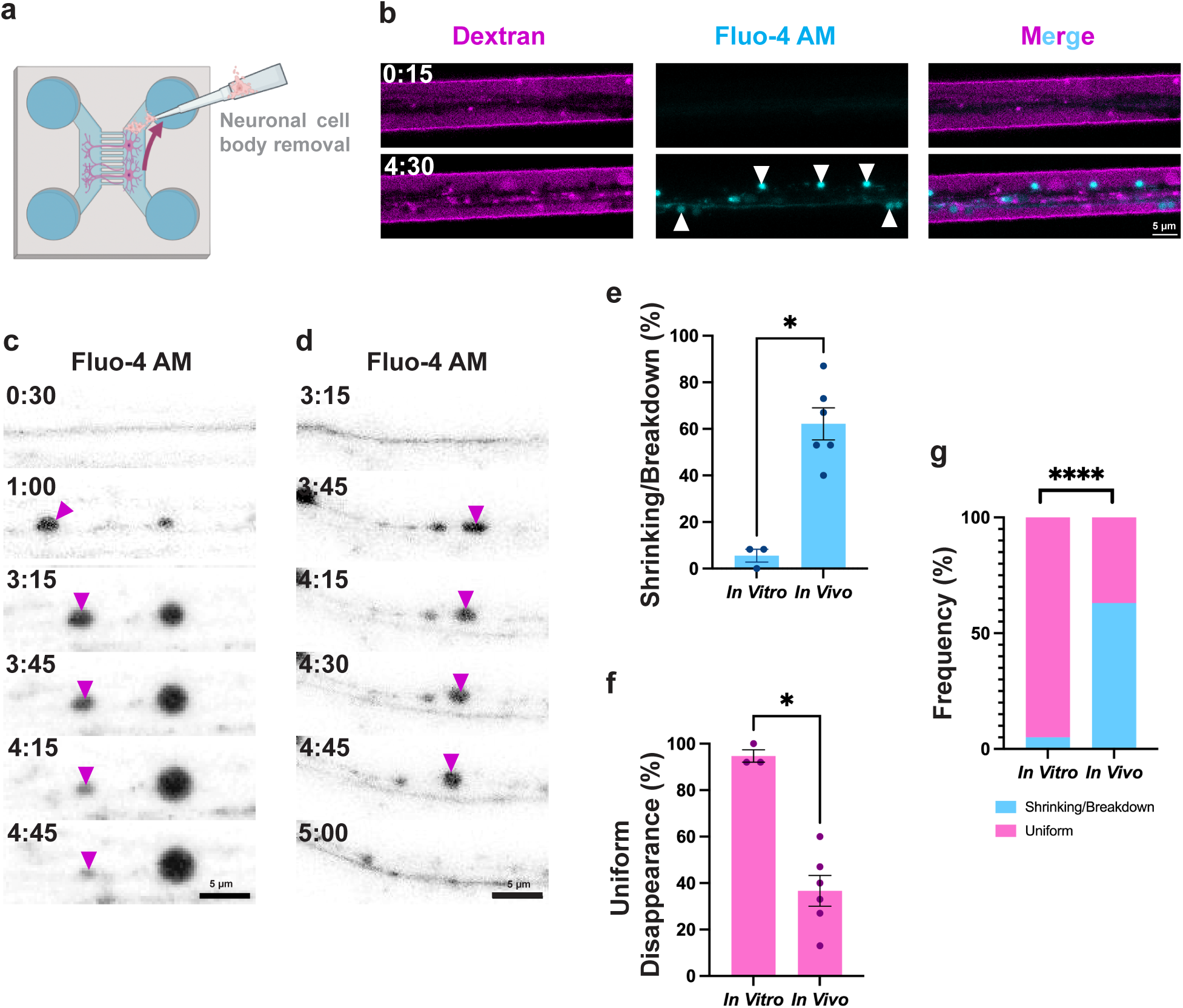
The extra-axonal environment is required for spheroid shrinking and breakdown. a) A model to study axonal spheroids *in vitro*: sympathetic neurons are plated in one chamber of a microfluidic device (MFD), then given 5-7 days for axons to grow across grooves into the opposite chamber. Cell bodies are aspirated, leaving behind the injured axons. Axons are imaged with time-lapse confocal microscopy using additional fluorescent dyes. b) Representative image of calcium-labeled (Fluo-4 AM) spheroids (arrowheads) with negative dextran labeling after injury *in vitro*. c) Fluo-4 AM-labeled spheroid (arrowheads) shrinking *in vitro*. d) A Fluo-4 AM-labeled spheroid undergoing uniform disappearance (arrowheads) *in vitro*. e) The frequencies of spheroid shrinking or breakdown *in vitro* and *in vivo* (N=3 MFDs and 6 larvae, p=0.0238, two-tailed Mann-Whitney U test). f) The frequencies of spheroid uniform disappearance *in vitro* and *in vivo* (N=3 MFDs and 6 larvae, p=0.0238, two-tailed Mann-Whitney U test). g) Relative frequencies of shrinking/breakdown and uniform disappearance *in vitro* and *in vivo* (N=3 MFDs and 6 larvae, p<0.0001, two-sided Fisher’s Exact Test).

Comparing *in vitro* and *in vivo* time-lapses, we found that spheroid fates differed between these conditions. *In vitro*, we primarily observed spheroids uniformly disappearing (95 ± 2.7%, Fig. 3d, f, g) with only 5.5 ± 2.8% of spheroids shrinking (Fig. 3c) and virtually no detectable spheroids breaking down (Fig. 3 e, g). *In vivo*, 37 ± 6.7% of spheroids disappeared uniformly, while 62 ± 6.9% shrank or broke down (Fig. 3e, f, g). These differences were statistically significant for both partial signal loss and uniform signal loss (Fig. 3e, f, g).

Collectively, these findings indicate that uniform disappearance can occur independently of the extra-axonal environment, consistent with membrane permeabilization and cytoplasmic content release as previously reported <u>[15]</u>. However, these data demonstrate that the extra-axonal environment is required for spheroid shrinking and breakdown.

### Schwann cells interact with axonal spheroids

We next considered which factors in the extra-axonal environment could give rise to different spheroid fates. Patterns of shrinking and breakdown are reminiscent of cell-cell interactions, such as dendritic bead shrinking by microglia after seizure <u>[34]</u> or cell-assisted disassembly in apoptotic body formation <u>[33]</u>. Therefore, we hypothesized that interactions with adjacent glial cells may underlie the differences in spheroid fates. Specifically, we chose to investigate SCs due to their known roles in responding to nerve injury and their proximity to spheroids <u>[20,23,24,35]</u>.

To characterize SC-spheroid proximity, we axotomized the pLLN of *sox10:mRFP;HuC:GCaMP6s* larvae, where *sox10* regulatory sequences drive expression of a membrane-tethered, red-fluorescent protein in SCs. After injury, we observed that SCs maintained contact with both axons and spheroids (Fig. 4a, b). When we quantified this contact as the percentage of spheroid surface area touching SCs, we found that spheroids had an average overlap ratio of 87 ± 0.79% (Fig. 4c), and 72% of spheroids had an overlap ratio of 0.95 or higher (Fig. 4d). From these data, we concluded that SCs are in contact with most axonal spheroids after injury.

**Figure 4:**
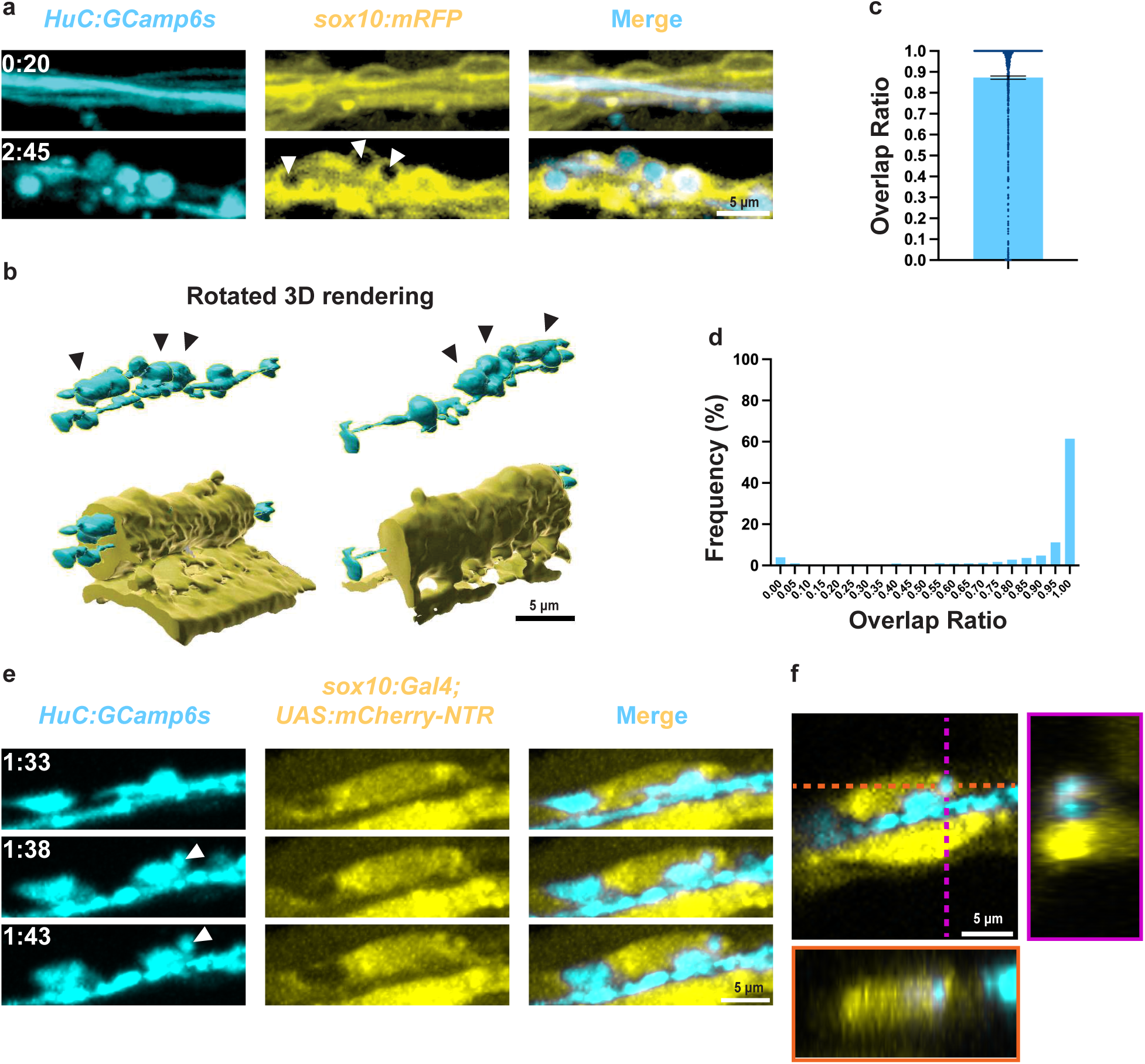
Schwann cells interact with axonal spheroids. a) Spheroids (*HuC:G-Camp6s*) localize within Schwann cells (*sox10:mRFP*, white arrowheads. Maximum z projection, top, and single z plane, bottom). b) 3D projection of a) showing spheroids (black arrowheads) within Schwann cell membranes. c) Percentages of spheroid surfaces covered by Schwann cell, or overlap ratio (N=1,067 spheroids from 6 fish, error bar = SEM). d) Histogram of individual spheroid overlap ratios in c). e) A spheroid (*HuC:G-Camp6s*, white arrowhead) is brought into the Schwann cell cytoplasm (*sox10:-Gal4;UAS:NTR-mCherry*). f) Single plane XY (square panel), YZ (magenta), and XZ (orange) cross sections of the spheroid in e inside Schwann cell cytoplasm.

Because *sox10:mRFP* signal is localized to the membrane, it was difficult to determine whether spheroids were in contact with SCs extracellularly or intracellularly. To determine whether SC-spheroid contact occurred intracellularly, we examined spheroid localization relative to SC cytosol. To do this, we imaged 3 dpf *sox10:Gal4;UAS:mCherry-NTR;HuC:GCaMP6s* larvae after pLLN axotomy. In these larvae, *sox10* regulatory sequences drive Gal4 expression. Gal4 binds to a UAS site to promote expression of a cytoplasmic mCherry protein fused to an inactive nitroreductase enzyme. With this cytoplasmic SC marker, we observed a subset of spheroids that moved off the axon to inside contiguous SC cytoplasm (Fig. 4e, f, Supplementary video 4). Taken together, we conclude that SCs are in contact with the vast majority of spheroids via their membranes and that this contact is not limited to the extracellular SC compartment, as at least a subset of these interactions appear to be intracellular within the SC.

### Spheroids expose phosphatidylserine

To investigate how SCs might interact with spheroids after injury, we examined a canonical “eat me” signal, phosphatidylserine (PS), which resides on the outer membrane leaflets of dying cells and facilitates interactions with phagocytic receptors <u>[36]</u>. We and others previously observed that spheroids expose PS *in vitro <u>[26,27]</u>*, and SCs engage in debris clearance after nerve injury through PS receptor MerTK <u>[24]</u>. Therefore, we hypothesized that PS may similarly be displayed on axonal spheroids during degeneration *in vivo*. To investigate this, we first verified PS exposure *in vitro* after axotomy using our *in vitro* model with negative dextran labeling and visualizing PS exposure with fluorescent annexin V. Because annexin V binds to exposed PS, its visualization indicates the presence of PS on outer membrane leaflets <u>[37]</u>. From these time-lapse images, we confirmed that spheroids expose PS after axotomy *in vitro* (Fig. 5a).

**Figure 5:**
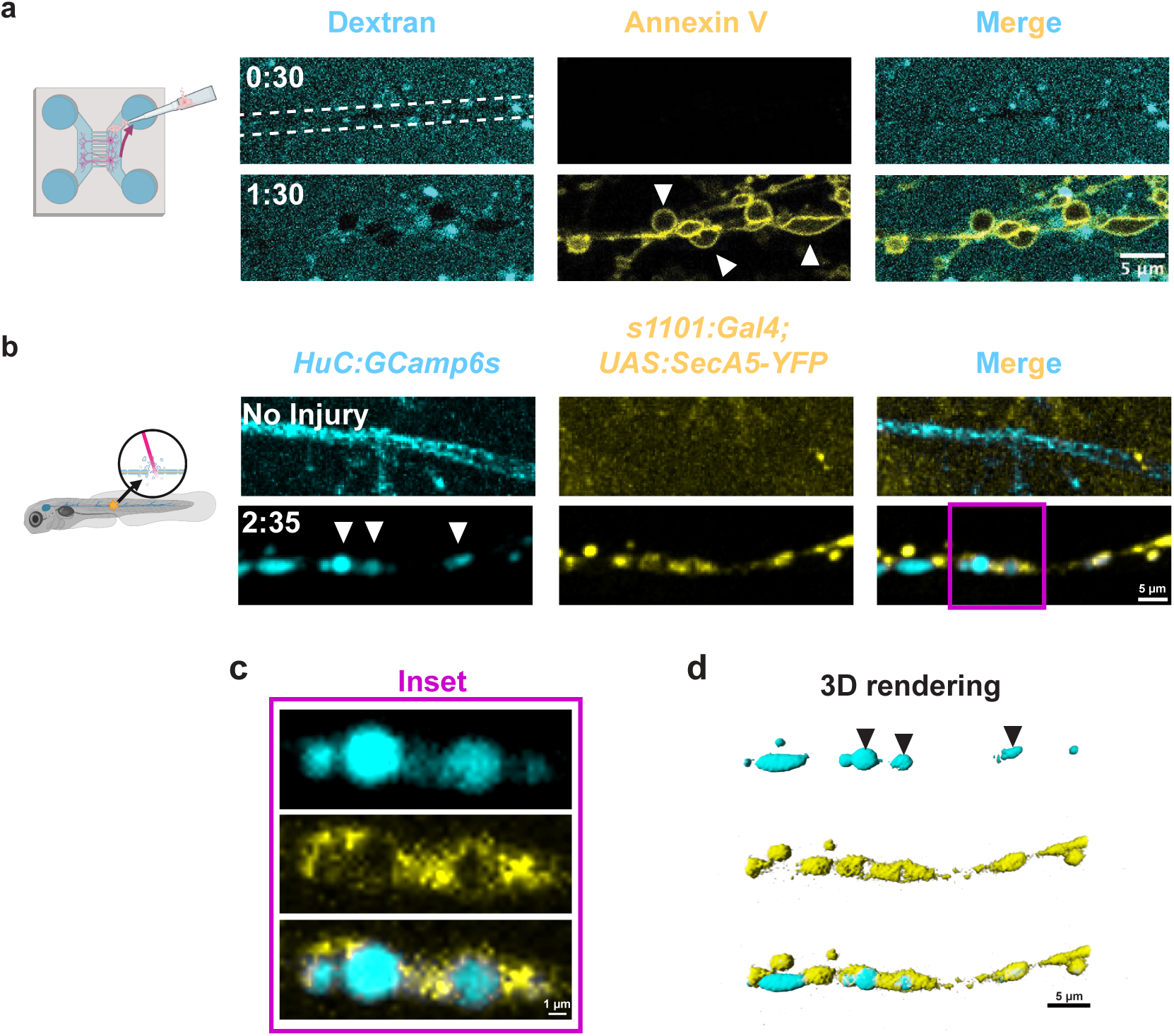
Spheroids expose phosphatidylserine. a) Axonal spheroids expose phosphatidylserine (PS, annexin V, white arrowheads) after *in vitro* axotomy, dashed lines indicate negative axon staining (dextran, N=3 MFDs). b) Spheroids (*HuC:GCaMP6s*) expose phosphatidylserine after axotomy *in vivo* (*s1101:Gal4;UAS:SecA5-YFP*, white arrowheads, inset showing single z slice, N=5 larvae). c) Inset from b. d) 3D rendering of b showing PS-exposing spheroids (black arrowheads).

To determine whether spheroids expose PS *in vivo*, we visualized PS exposure on spheroids using *HuC:jRGECO1b;s1101:Gal4;UAS:SecA5-YFP* larvae. In these larvae, all post-mitotic neurons express jRGECO1b, a red fluorescent calcium indicator <u>[38]</u>. The *s1101* enhancer drives neuronal Gal4 expression <u>[39]</u>, which binds a UAS sequence and promotes downstream expression of secreted annexin V fused to a yellow fluorescent protein <u>[40]</u>. We observed PS in beads-on-a-string patterns after injury, leading us to conclude that spheroids expose PS *in vivo* (Fig. 5b; Supplementary video 5). However, PS and calcium rarely labeled the same spheroids, consistent with previous findings that axonal calcium influx is not required for PS exposure <u>[41]</u>. In instances where calcium and PS signals did coincide, PS signal appeared unevenly distributed around the spheroids, rather than in the even, consistent rings observed *in vitro* (Fig. 5c, d). Despite the different PS morphologies across *in vitro* and *in vivo* conditions, PS was present on spheroids in both models, demonstrating that spheroids present an “eat me” signal known to interact with SCs after injury.

### Schwann cells induce shrinking and breakdown in spheroids

Given that the extra-axonal environment alters spheroid fates and SCs interact with axonal spheroids, we next investigated whether SCs alter spheroid fates. We employed a *sox10:Gal4;UAS:mCherry-NTR* transgenic line for SC-specific ablation. In this system, the nitroreductase (NTR) enzyme fused to mCherry produces reactive oxygen species in the presence of Ronidazole (RDZ), causing *sox10-*positive cell death <u>[42]</u>. We incubated 2 dpf larvae in 4 mM RDZ or vehicle (0.003% phenylthiourea in egg water), then axotomized the pLLNs and imaged them at 3 dpf.

We first assessed the effects of SC ablation on spheroid quantities over time after axotomy. We observed no difference in spheroid quantities between control and SC ablation conditions from 0 to 120 minutes post-injury. However, after 120 minutes, we observed more spheroids persisting up to 300 minutes post-injury in control conditions compared to SC ablation conditions (Fig. 6a, b, Supplementary video 6, Supplementary video 7). We determined that differences after 120 minutes were statistically significant using Poisson mixed-effect modeling with restricted cubic splines (Fig. 6b; Supplementary Materials 2). Additionally, we found that average spheroid lifetimes were significantly shorter in SC ablation conditions compared to vehicle control (VC, Fig. 6c, d). These findings demonstrate that decreased spheroid quantity observed in SC ablation conditions was due to faster spheroid loss rather than reduced spheroid formation.

**Figure 6:**
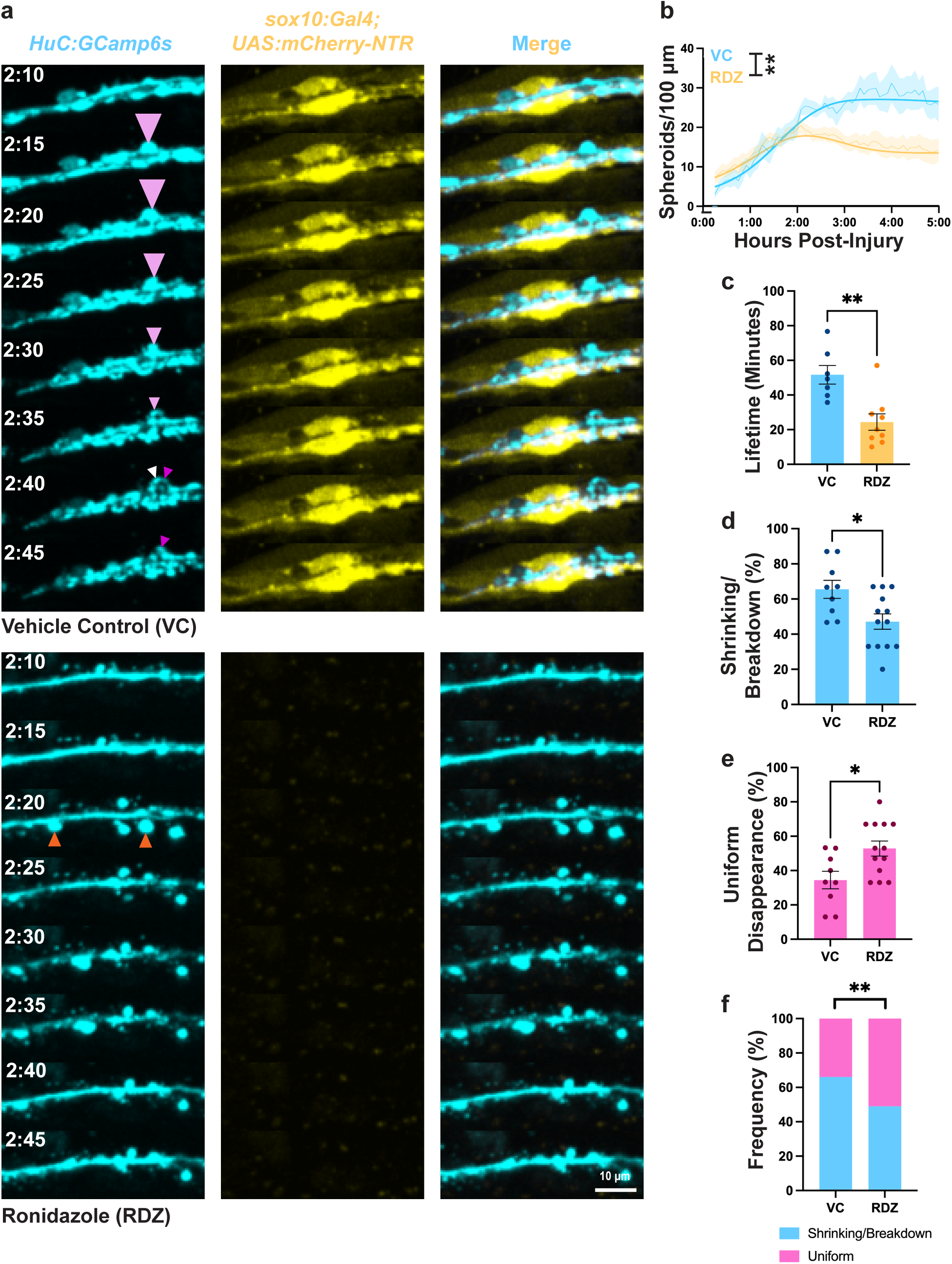
Schwann cells induce shrinking and breakdown in spheroids. a) pLLNs (*HuC:GCamp6s*) from larvae treated 2-3dpf with 4mM Ronidazole (RDZ) or vehicle control (VC) to ablate Schwann cells (SCs, *sox10:Gal4; UAS:NTR-mCherry*). Spheroids frequently undergo shrinking (pink arrowheads) and breakdown (pink to white and magenta arrowheads) in VC treatment, and uniform disappearance (orange arrowheads) in RDZ treatment. b) Spheroid quantities over time (thin lines) predict significantly different values (thick lines) between treatment groups after 120 minutes (N=14 RDZ, 6 VC, p<0.005, Poisson mixed-effects model, Supplementary Materials 2). c) Average spheroid lifetimes (N=7 VC, 8 RDZ larvae, p=0.0033, two-tailed Mann-Whitney U Test). d) Frequences of spheroid shrinking or breakdown (N=9 VC, 13 RDZ, p=0.0335, two-tailed Mann-Whitney U Test). e) Relative frequencies of shrinking/breakdown and uniform disappearance (N=9 VC, 13 RDZ, p=0.0044, two-sided Fisher’s Exact Test).

Intriguingly, spheroids present at later times in control conditions appeared to be primarily broken-down spheroids with persisting signal (Supplementary video 6, Supplementary video 7). To verify this, we examined the frequency of spheroid disappearance patterns in each condition. We found that SC ablation led to a significant decrease in spheroid shrinking and breakdown, accompanied by a significant increase in uniform disappearance (Fig. 6d).

Collectively, these findings demonstrate that SCs alter spheroid dynamics by promoting shrinking and breakdown of spheroids. In the presence of SCs, spheroids tend to persist longer and undergo shrinking and breakdown, whereas in their absence, spheroids are more likely to undergo rapid, uniform disappearance. Figure 7 presents a graphical model summarizing these results, illustrating the crucial role of SCs in modulating spheroid fate and longevity.

**Figure 7:**
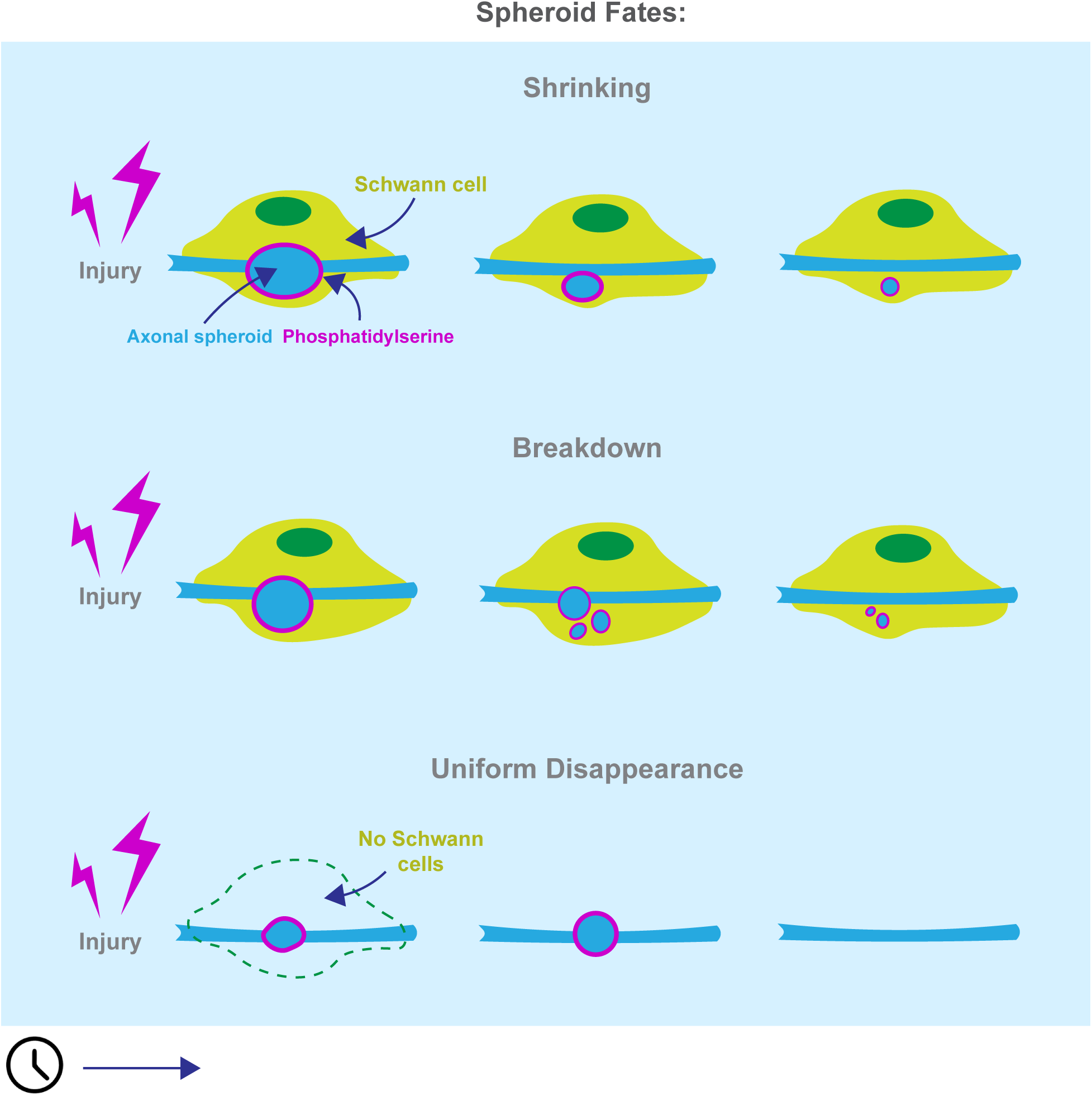
Working model for Schwann cell effects on spheroid fates. Following injury, spheroids are more likely to undergo shrinking and breakdown when Schwann cells (SCs) are present on degenerating axons. When Schwann cells are absent, spheroids are more likely to undergo uniform disappearance. Spheroids expose phosphati-dylserine, which could facilitate interactions between spheroids and SCs.

## Discussion

In this study using an *in vivo* model to investigate axonal spheroid dynamics, we observed that spheroids are present before, during, and after axon fragmentation and that individual spheroids vary in onset and duration times. When spheroids disappear, they shrink, break down into puncta, or vanish uniformly without any apparent morphological changes. The absence of spheroid shrinking and breakdown *in vitro* suggests that these fates require interactions with the extra-axonal environment. In keeping with this, SCs enwrap and internalize spheroids, which expose PS during degeneration, and SCs are required for spheroid shrinking and breakdown *in vivo*.

Previous studies have characterized axonal spheroid formation *in vitro* and *ex vivo <u>[11,13,14]</u>*, focusing on periods before and during degeneration. We extend these characterizations by examining spheroid dynamics in an *in vivo* model and track their progression from formation to disappearance. We observed that individual spheroids exhibit variable lifetimes, persisting both before and after axon fragmentation (Fig. 1d, e). Whether these differences correspond to distinct functional outcomes remains unknown but might suggest a link between spheroid fate and function <u>[15,19]</u>. A minority of spheroids persisted into the regenerative window, up to 300 minutes post-injury (Fig. 1e), raising questions about their long-term impact on axon integrity and potential influence on regeneration.

We identified three patterns of spheroid disappearance after peripheral nerve injury *in vivo*: shrinking, breakdown, and uniform disappearance (Fig. 2). Spheroid shrinking has been observed in Alzheimer’s Disease <u>[43]</u>. Extending this phenomenon to peripheral nerve injury suggests conservation across central and peripheral nervous systems, cell types, and degenerative etiologies. Uniform spheroid disappearance has also been characterized *in vitro*, attributed to membrane permeabilization <u>[15]</u>. Before our study, spheroids breaking down into puncta had not been reported, possibly because prior research utilized chronic degeneration models with longer imaging intervals that missed rapid breakdown events <u>[19,43]</u>, or because they were conducted *in vitro* where we did not observe this fate <u>[15,26]</u>. If breakdown into puncta is conserved across degenerative triggers in the same way that shrinking is, it may represent a widespread spheroid fate. Importantly, our characterization of spheroid fates is based on cytoplasmic morphology, which may limit interpretations regarding membrane integrity and overall structure. Future research describing these fates with respect to membrane dynamics would shed light on each fate’s corresponding cellular processes.

The prevalence of spheroid shrinking and breakdown *in vivo*, but not *in vitro* or after SC ablation (Fig. 2, Fig. 3, Fig. 6), suggests that interactions with extra-axonal factors such as glial cells contribute to these fates. The exact cellular processes underlying these fates remain unclear. Possible mechanisms include trogocytosis or calcium buffering in the case of shrinking spheroids <u>[44]</u>, spheroid disassembly into apoptotic bodies or signal quenching around membrane-bound organelles for spheroid breakdown <u>[45]</u>, and phagocytic engulfment or rupture for uniform disappearance <u>[15]</u>. Moreover, the incomplete deficit in shrinking and breakdown following SC ablation suggests that other cell types may also facilitate these spheroid fates (Fig. 6). This is further supported by spheroid shrinking in the CNS where SCs are absent <u>[19,34,43]</u>. Therefore, while these studies make headway in characterizing spheroid fates, future work to understand their cellular and molecular underpinnings remains to be done.

To further elucidate the cellular processes mediating spheroid fates, we characterized the interactions between spheroids and SCs (Fig. 4). We found that SCs contact most spheroids with their membranes and that some spheroids are within SC cytoplasm (Fig. 4, Supplementary video 4), indicating active interactions. Additionally, we showed that spheroids expose PS after peripheral nerve injury *in vivo* (Fig. 5), complementing prior descriptions of PS exposure after trophic deprivation and after axotomy *in vitro <u>[26,27]</u>*. *In vitro,* we observed even, ring-like PS labeling around spheroids. In contrast, *in vivo*, PS-binding annexin V was often aggregated into puncta around calcium-labeled spheroids (Fig. 5, Supplementary video 5). These distinct patterns may result from methodological differences; *in vitro*, abundant annexin V in the extracellular medium labels PS uniformly, whereas *in vivo*, the secreted annexin V protein may have limited availability due to synthesis rates, diffusion constraints, or calcium buffering <u>[37]</u>. Either way, PS was present on spheroids, an interesting finding given that PS interactions are conserved across a breadth of unprofessional and professional phagocytes not limited to SCs <u>[46,47]</u>.

In conclusion, our work provides an *in vivo* characterization of spheroid dynamics from formation to disappearance, identifying three predominant fates: shrinking, breakdown, and uniform disappearance. It examines cellular mechanisms by which the extra-axonal environment could influence spheroid fates, establishing one cell type, SCs, that interact with spheroids and contribute to spheroid shrinking and breakdown. Collectively, our findings bridge the gap in understanding spheroid dynamics within the extra-axonal environment, establish that axonal spheroid fates are regulated in the native nerve environment, and highlight SCs as a key cellular mechanism influencing spheroid outcomes. This work paves the way for future research to elucidate how spheroid regulation and dysregulation contribute to degenerative processes in development and disease, ultimately identifying new targets for therapeutic intervention.

## Methods

### Animals

All mouse studies complied with the policies of the Association for Assessment and Accreditation of Laboratory Animal Care International (AAALAC) and were approved by the University of Virginia Institutional Animal Care and Use Committee (IACUC). All mice were on a C57BL/6 background obtained from Jackson Laboratory. Both male and female mice were used in all experiments.

All zebrafish studies were approved by the University of Virginia IACUC. Embryos were produced by pairwise natural matings, cleaned daily, and maintained at 28.5 °C in egg water. At 1 day post-fertilization (dpf), embryos were treated with 0.0003% phenylthiourea (PTU) to prevent pigmentation. At 2 or 3 dpf, embryos were screened for transgenes of interest using an epifluorescence microscope. Larvae were imaged for experiments at 3 dpf, prior to sexual differentiation <u>[48]</u>.

**Table 1:**
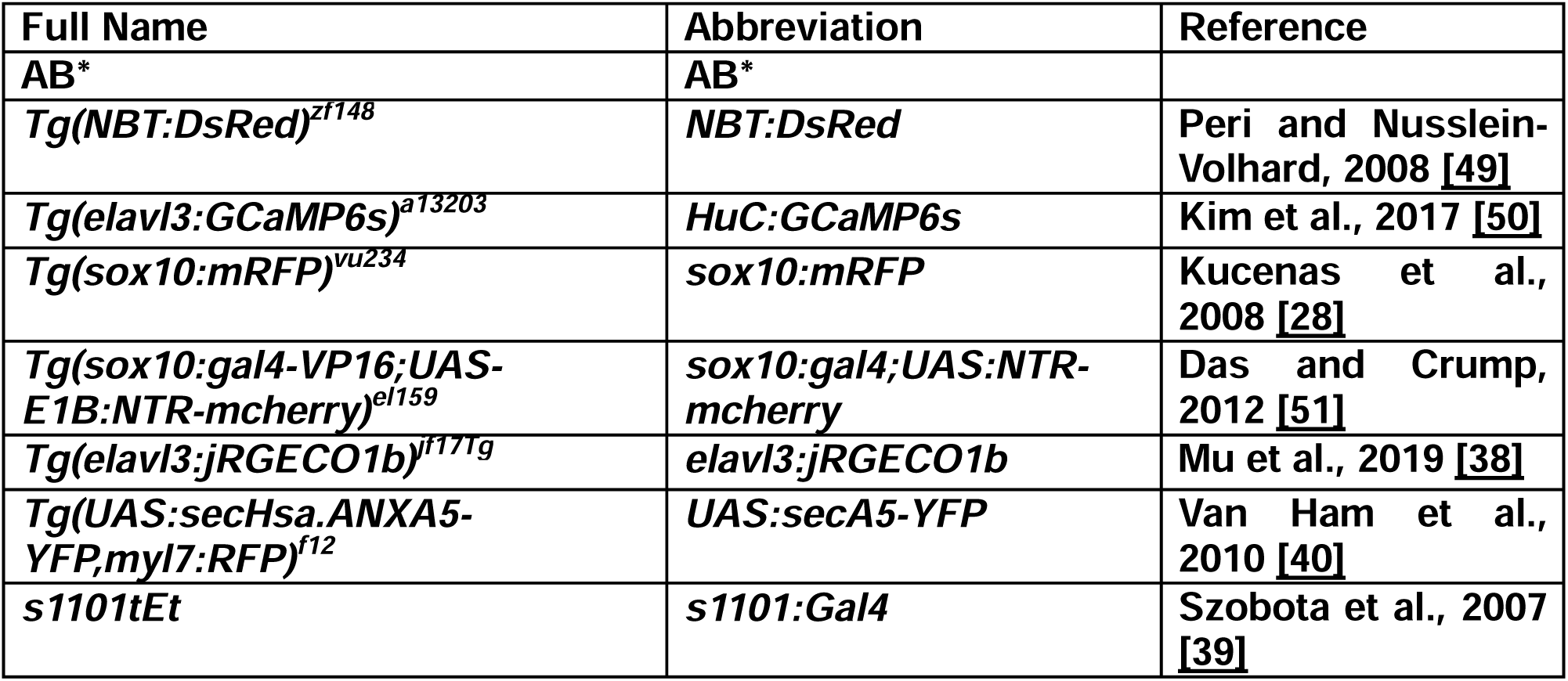
Transgenic Zebrafish Lines Used in This Study.

### Primary Superior Cervical Ganglion Cultures

Superior cervical ganglion (SCG) neurons were harvested from postnatal day 0–2 C57BL/6 mice as previously described <u>[52]</u>. SCGs were dissociated in a solution containing 0.01 g/mL bovine serum albumin, deoxyribonuclease I, 0.4 mg/mL hyaluronidase, and 4 mg/mL collagenase type II in DMEM for 15 minutes at 37 °C with 10% CO_₂_. The ganglia were then incubated in 2.5% trypsin in DMEM for 20 minutes at 37 °C with 10% CO_₂_. Cells were triturated and centrifuged at 300 × g for 5 minutes at 4 °C. The supernatant was aspirated, and cells were resuspended in 3.5 µL of SCG medium per ganglion harvested. The SCG medium consisted of DMEM/F12 supplemented with 10% fetal bovine serum, penicillin/streptomycin (1 U/mL), and nerve growth factor (45 ng/mL, isolated from mouse salivary glands). A volume of 3.2 µL of the SCG suspension was plated into microfluidic devices (MFDs), manufactured as previously described <u>[53]</u> and adhered to coverslips coated with 50 µg/mL poly-D-lysine and 1 µg/mL laminin. Cells were allowed to adhere for 20 minutes at 37 °C. SCG medium (100 µL) was added to each distal axon chamber well, and 125 µL to each cell body well. Cultures were maintained at 37 °C with 10% CO_₂_. Neurons were purified by treating with cytosine arabinofuranoside (AraC) from days 1 to 3 in vitro (DIV 1–3). Axon extension into the adjacent chamber occurred between DIV 5–7.

### *In Vitro* Axotomy Live Imaging Experiments

At DIV 5–7, SCG medium was removed and replaced with phenol red–free DMEM/F12 medium containing either Fluo-4 AM (1 µM), Incucyte Annexin V Dye for Apoptosis, NIR (1:200; 4768, Sartorius), or 3,000 MW Cascade Blue Dextran (100 µM), depending on the experiment. Cultures were incubated for 30 minutes at 37 °C with 10% CO_₂_. The medium in the cell body wells was removed and set aside, and neuronal cell bodies were then aspirated to induce axotomy <u>[53]</u>, confirmed via bright-field microscopy. Axons were imaged in the grooves connecting the MFD chambers using a Zeiss LSM 980 confocal microscope with a 63× oil immersion objective. Images were captured as 1 µm slice z-stacks from 15 to 300 minutes post-injury at 15-minute intervals.

### *In Vivo* Axotomy Live Imaging Experiments

At 3 dpf, zebrafish larvae were anesthetized in Tricaine and mounted in 0.8% low– melting point agarose in glass-bottomed Petri dishes. PTU egg water was added over the larvae for imaging on a Zeiss AxioObserver Z1 microscope configured with Quorum WaveFX-XI software, an Andor CSU-W1 spinning disk confocal system, or a Dragonfly spinning disk confocal system. All image acquisition was performed with a 40× water immersion objective. Axotomies were performed around the 18th somite with a nitrogen dye (435 nm)–pumped MicroPoint laser. Ablation regions of interest (ROIs) approximately 5 µm in diameter were created on the pLLN around the 18th somite using the 40× objective. Complete pLLN transections were confirmed by observing acute axon degeneration <u>[2]</u> and the absence of intact axon signal across the injury site. Images were centered 100 µm distal to the injury site, and 1 µm slice z-stacks (22–29 µm depth) were acquired for 300 minutes at intervals of 1.5, 3, or 5 minutes, depending on the experiment

### Ronidazole Treatment

For Schwann cell ablation, larvae expressing *HuC:GCaMP6s* and *sox10:Gal4; UAS:NTR-mCherry* were incubated in the dark at 28.5 °C from 2 dpf to 3 dpf in 4 mM Ronidazole (RDZ) in PTU egg water or vehicle control.

### Image Processing

*In vivo* time-lapse images were processed using Imaris software. Videos were oriented so that the anterior/posterior axis was left/right, and supplementary videos were oriented so that the dorsal/ventral axis was top/bottom. Unless otherwise specified, videos and images are presented as maximum intensity z-projections. Brightness and contrast were manually adjusted for visibility. Regions of interest (ROIs) for quantifications were created by cropping videos to the region 150 to 250 µm distal to the injury site. Three-dimensional surface renderings were made using the Bitplane Imaris surface creation module.

*In vitro* time-lapse images were processed in FIJI (ImageJ). Fluo-4 AM and annexin V images were presented as maximum intensity z-projections, and dextran images were presented as single z-planes 1 µm thick. Brightness and contrast were manually adjusted for visibility.

### Quantification and Statistical Analysis

We conducted quantitative analyses to assess spheroid dynamics and interactions using Imaris, FIJI, GraphPad Prism, R, and R Studio software. Quantifications were performed on time-lapses processed as described above using frames acquired at times closest to the intervals of interest.

- **Spheroid Counts Over Time:** Spheroids were detected using machine learning segmentation in Imaris from time-lapse videos cropped to the ROI. Counts were recorded at 5-minute intervals post-injury. Comparisons between treatments (i.e., with and without RDZ) were analyzed using Poisson mixed-effect models with restricted cubic splines (Supplementary Materials 2).
- **Validation of Automated Detection:** Automated spheroid detection using machine learning segmentation in Imaris was validated against manual counts by calculating precision and sensitivity (Supplementary Fig. 2). Linear mixed-effects modeling assessed the relationship between automated and manual counts (Supplementary Materials 1).
- **Time to Nerve Perforation:** The time at which the last intact axon fragmented in each time-lapse video was recorded and plotted in GraphPad Prism.
- **Spheroid Lifetimes:** Spheroid formation and disappearance times were recorded to determine lifetimes, defined as the interval between formation and disappearance. At least 2 spheroids from at least 4 different calcium transients were quantified in each larva. Spheroids already present at 15 minutes were excluded from analyses. Lifetimes were binned into 15-minute intervals.
- **Spheroid Fates *In Vivo:*** Fates were categorized as shrinking/breakdown or uniform disappearance based on morphological changes. Analyses included at least 5 spheroids from at least 3 calcium transients per larva.
- **Comparison Between *In Vitro* and *In Vivo* Fates:** To compare spheroid fates between models, time-lapse videos were analyzed at equivalent intervals (15 minutes). *In vitro,* spheroid fates were analyzed at 15 minute intervals. Spheroids that did not disappear by the end of the time lapse were excluded from analyses. At least 10 spheroids from at least 3 different calcium transients were categorized *in vitro. In vivo,* time-lapses were edited 15 minute intervals. At least 15 spheroids from at least 4 calcium transients were quantified.
- **Comparison Between Spheroid Fates +/- Ronidazole:** Spheroid fates were categorized by a blinded reviewer as broken down/shrinking or not. Spheroids that did not break down or shrink were counted as uniform disappearances. Analyses included at least 5 spheroids from at least 3 calcium transients per larva.
- **Spheroid–Schwann Cell Contact Analysis:** Schwann cell surfaces and spheroid surfaces were defined with Imaris surface creation. The overlap ratio, defined as the percentage of spheroid surface area in contact with Schwann cells, was calculated for each spheroid.
- **Data Collection and Statistical Methods:** Data were collected from at least three independent experiments for both *in vivo* (N=3 clutches) and *in vitro* (N=3 litters/MFDs) studies. For comparisons between conditions (e.g., with and without RDZ) a blinded reviewer conducted quantifications. Statistical analyses were performed using GraphPad Prism and R, with specific tests detailed in the figure legends and supplemental materials. Data are presented as means ± standard error of the mean (SEM), and significance was set at *p* < 0.05.

### Graphics

Graphical illustrations were created using Biorender and Adobe Illustrator.

## Acknowledgements

We thank Dr. Anthony Spano for technical assistance, and Lori Tocke and The University of Virginia Gilmer Hall vivarium staff for assistance with animal care. We thank Dr. Yu Yong for her foundational work establishing the knowledge and protocols upon which this study builds, and Riya Verma for help with earlier experiments leading up to these studies. We thank Dr. Ammasi Periasamy and acknowledge the Keck Center for Cellular Imaging for the usage of the Zeiss 880/980 microscopy system. [PI: AP; NIH-0D0-25156]. We thank Dr. Christopher Hughes and the James Madison University physics department for their assistance manufacturing microfluidic devices. We also thank Dr. Amber English for her guidance in implementing the Bitplane - Imaris FIJI plugin machine learning segmentation algorithm for automated spheroid detection.

This work was supported by the National Institutes of Health [Grant 5R01NS091617, Grant 1R21NS124164, Grant 5R01NS107525] and The Robert R. Wagner Fellowship.

## Author contributions

S.H.C. proposed and designed these studies with feedback from E.S., S.K., and C.D.D. S.H.C. carried out these studies with help from T.V., C.K.A., B.M., and H.K. on cell culture establishment and care, zebrafish embryo care and transgenic embryo screening, and with help from H.K. on *in vitro* image acquisitions. S.H.C. interpreted all of the data with major contributions from C.K.A., T.V., E.S., H.K., and B.M. on blinded image analyses, and from B.M. and M.J. on data analysis in R. M.J. performed Poisson mixed-effect modeling and statistical oversight. S.H.C. performed other statistical analyses. S.H.C. prepared the figures and text for this manuscript with substantial edits from and feedback from S.K. and C.D.D.

## Data availability statement

The raw datasets used and/or analyzed in the current study are available from the corresponding authors upon reasonable request.

## Declaration of generative AI and AI-assisted technologies in the writing process

During the preparation of this work S.H.C. used ChatGPT in order to identify prospective appropriate statistical analyses and write scripts for data analysis in R. C.D. used ChatGPT and Anthropic to edit manuscript text for grammar and overall structure. After using this tool, the authors reviewed and edited the content as needed and take full responsibility for the content of the published article.

## References

1. Landowski, L.M., Dyck, P.J.B., Engelstad, J., and Taylor, B.V. (2016). Axonopathy in peripheral neuropathies: Mechanisms and therapeutic approaches for regeneration. J. Chem. Neuroanat. 76, 19–27.

2. Wang, J.T., Medress, Z.A., and Barres, B.A. (2012). Axon degeneration: molecular mechanisms of a self-destruction pathway. J. Cell Biol. 196, 7–18.

3. Cavanagh, J.B. (1964). The significance of the “dying back” process in experimental and human neurological disease. Int. Rev. Exp. Pathol. 3, 219–267.

4. Sagot, Y., Dubois-Dauphin, M., Tan, S.A., de Bilbao, F., Aebischer, P., Martinou, J.C., and Kato, A.C. (1995). Bcl-2 overexpression prevents motoneuron cell body loss but not axonal degeneration in a mouse model of a neurodegenerative disease. J. Neurosci. 15, 7727–7733.

5. Samuel, M.A., Zhang, Y., Meister, M., and Sanes, J.R. (2011). Age-related alterations in neurons of the mouse retina. J. Neurosci. 31, 16033–16044.

6. Saxena, S., and Caroni, P. (2007). Mechanisms of axon degeneration: from development to disease. Prog. Neurobiol. 83, 174–191.

7. Schoenmann, Z., Assa-Kunik, E., Tiomny, S., Minis, A., Haklai-Topper, L., Arama, E., and Yaron, A. (2010). Axonal degeneration is regulated by the apoptotic machinery or a NAD+-sensitive pathway in insects and mammals. J. Neurosci. 30, 6375–6386.

8. Simon, D.J., Weimer, R.M., McLaughlin, T., Kallop, D., Stanger, K., Yang, J., O’Leary, D.D.M., Hannoush, R.N., and Tessier-Lavigne, M. (2012). A caspase cascade regulating developmental axon degeneration. J. Neurosci. 32, 17540–17553.

9. Osterloh, J.M., Yang, J., Rooney, T.M., Fox, A.N., Adalbert, R., Powell, E.H., Sheehan, A.E., Avery, M.A., Hackett, R., Logan, M.A., et al. (2012). dSarm/Sarm1 is required for activation of an injury-induced axon death pathway. Science 337, 481–484.

10. Geden, M.J., and Deshmukh, M. (2016). Axon degeneration: context defines distinct pathways. Curr. Opin. Neurobiol. 39, 108–115.

11. Yong, Y., Hunter-Chang, S., Stepanova, E., and Deppmann, C. (2021). Axonal spheroids in neurodegeneration. Mol. Cell. Neurosci. 117, 103679.

12. Ramón y Cajal, S. Degeneration and regeneration of the nervous system. Clarendon Press, 1928.

13. Barsukova, A.G., Forte, M., and Bourdette, D. (2012). Focal increases of axoplasmic Ca2+, aggregation of sodium-calcium exchanger, N-type Ca2+ channel, and actin define the sites of spheroids in axons undergoing oxidative stress. J. Neurosci. 32, 12028–12037.

14. Beirowski, B., Nógrádi, A., Babetto, E., Garcia-Alias, G., and Coleman, M.P. (2010). Mechanisms of axonal spheroid formation in central nervous system Wallerian degeneration. J. Neuropathol. Exp. Neurol. 69, 455–472.

15. Yong, Y., Gamage, K., Cushman, C., Spano, A., and Deppmann, C. (2020). Regulation of degenerative spheroids after injury. Sci. Rep. 10, 15472.

16. Ohgami, T., Kitamoto, T., and Tateishi, J. (1992). Alzheimer’s amyloid precursor protein accumulates within axonal swellings in human brain lesions. Neurosci. Lett. 136, 75–78.

17. Newell, K.L., Boyer, P., Gomez-Tortosa, E., Hobbs, W., Hedley-Whyte, E.T., Vonsattel, J.P., and Hyman, B.T. (1999). Alpha-synuclein immunoreactivity is present in axonal swellings in neuroaxonal dystrophy and acute traumatic brain injury. J. Neuropathol. Exp. Neurol. 58, 1263–1268.

18. Ferreirinha, F., Quattrini, A., Pirozzi, M., Valsecchi, V., Dina, G., Broccoli, V., Auricchio, A., Piemonte, F., Tozzi, G., Gaeta, L., et al. (2004). Axonal degeneration in paraplegin-deficient mice is associated with abnormal mitochondria and impairment of axonal transport. J. Clin. Invest. 113, 231–242.

19. Yuan, P., Zhang, M., Tong, L., Morse, T.M., McDougal, R.A., Ding, H., Chan, D., Cai, Y., and Grutzendler, J. (2022). PLD3 affects axonal spheroids and network defects in Alzheimer’s disease. Nature 612, 328–337.

20. Salzer, J.L. (2012). Axonal regulation of Schwann cell ensheathment and myelination. J. Peripher. Nerv. Syst. 17 Suppl 3, 14–19.

21. Heredia, D.J., De Angeli, C., Fedi, C., and Gould, T.W. (2020). Calcium Signaling in Schwann cells. Neurosci. Lett. 729, 134959.

22. Bouçanova, F., and Chrast, R. (2020). Metabolic interaction between schwann cells and axons under physiological and disease conditions. Front. Cell. Neurosci. 14, 148.

23. Babetto, E., Wong, K.M., and Beirowski, B. (2020). A glycolytic shift in Schwann cells supports injured axons. Nat. Neurosci. 23, 1215–1228.

24. Brosius Lutz, A., Chung, W.-S., Sloan, S.A., Carson, G.A., Zhou, L., Lovelett, E., Posada, S., Zuchero, J.B., and Barres, B.A. (2017). Schwann cells use TAM receptor-mediated phagocytosis in addition to autophagy to clear myelin in a mouse model of nerve injury. Proc Natl Acad Sci USA 114, E8072–E8080.

25. Fernandez-Valle, C., Bunge, R.P., and Bunge, M.B. (1995). Schwann cells degrade myelin and proliferate in the absence of macrophages: evidence from in vitro studies of Wallerian degeneration. J. Neurocytol. 24, 667–679.

26. Yong, Y., Gamage, K., Cheng, I., Barford, K., Spano, A., Winckler, B., and Deppmann, C. (2019). p75NTR and DR6 Regulate Distinct Phases of Axon Degeneration Demarcated by Spheroid Rupture. J. Neurosci. 39, 9503–9520.

27. Ko, K.W., Devault, L., Sasaki, Y., Milbrandt, J., and DiAntonio, A. (2021). Live imaging reveals the cellular events downstream of SARM1 activation. eLife 10.

28. Kucenas, S., Takada, N., Park, H.-C., Woodruff, E., Broadie, K., and Appel, B. (2008). CNS-derived glia ensheath peripheral nerves and mediate motor root development. Nat. Neurosci. 11, 143–151.

29. Limbach, L.E., Penick, R.L., Casseday, R.S., Hyland, M.A., Pontillo, E.A., Ayele, A.N., Pitts, K.M., Ackerman, S.D., Harty, B.L., Herbert, A.L., et al. (2022). Peripheral nerve development in zebrafish requires muscle patterning by tcf15/paraxis. Dev. Biol. 490, 37–49.

30. Lyons, D.A., and Talbot, W.S. (2014). Glial cell development and function in zebrafish. Cold Spring Harb. Perspect. Biol. 7, a020586.

31. Nazareth, L., St John, J., Murtaza, M., and Ekberg, J. (2021). Phagocytosis by peripheral glia: importance for nervous system functions and implications in injury and disease. Front. Cell Dev. Biol. 9, 660259.

32. Galloway, D.A., Phillips, A.E.M., Owen, D.R.J., and Moore, C.S. (2019). Phagocytosis in the brain: homeostasis and disease. Front. Immunol. 10, 790.

33. Atkin-Smith, G.K., and Poon, I.K.H. (2017). Disassembly of the dying: mechanisms and functions. Trends Cell Biol. 27, 151–162.

34. Eyo, U.B., Haruwaka, K., Mo, M., Campos-Salazar, A.B., Wang, L., Speros, X.S., Sabu, S., Xu, P., and Wu, L.-J. (2021). Microglia provide structural resolution to injured dendrites after severe seizures. Cell Rep. 35, 109080.

35. Vaquié, A., Sauvain, A., Duman, M., Nocera, G., Egger, B., Meyenhofer, F., Falquet, L., Bartesaghi, L., Chrast, R., Lamy, C.M., et al. (2019). Injured axons instruct schwann cells to build constricting actin spheres to accelerate axonal disintegration. Cell Rep. 27, 3152–3166.e7.

36. Segawa, K., and Nagata, S. (2015). An apoptotic “eat me” signal: phosphatidylserine exposure. Trends Cell Biol. 25, 639–650.

37. Logue, S.E., Elgendy, M., and Martin, S.J. (2009). Expression, purification and use of recombinant annexin V for the detection of apoptotic cells. Nat. Protoc. 4, 1383–1395.

38. Mu, Y., Bennett, D.V., Rubinov, M., Narayan, S., Yang, C.-T., Tanimoto, M., Mensh, B.D., Looger, L.L., and Ahrens, M.B. (2019). Glia Accumulate Evidence that Actions Are Futile and Suppress Unsuccessful Behavior. Cell 178, 27–43.e19.

39. Szobota, S., Gorostiza, P., Del Bene, F., Wyart, C., Fortin, D.L., Kolstad, K.D., Tulyathan, O., Volgraf, M., Numano, R., Aaron, H.L., et al. (2007). Remote control of neuronal activity with a light-gated glutamate receptor. Neuron 54, 535–545.

40. van Ham, T.J., Mapes, J., Kokel, D., and Peterson, R.T. (2010). Live imaging of apoptotic cells in zebrafish. FASEB J. 24, 4336–4342.

41. Shacham-Silverberg, V., Sar Shalom, H., Goldner, R., Golan-Vaishenker, Y., Gurwicz, N., Gokhman, I., and Yaron, A. (2018). Phosphatidylserine is a marker for axonal debris engulfment but its exposure can be decoupled from degeneration. Cell Death Dis. 9, 1116.

42. Curado, S., Stainier, D.Y.R., and Anderson, R.M. (2008). Nitroreductase-mediated cell/tissue ablation in zebrafish: a spatially and temporally controlled ablation method with applications in developmental and regeneration studies. Nat. Protoc. 3, 948–954.

43. Blazquez-Llorca, L., Valero-Freitag, S., Rodrigues, E.F., Merchán-Pérez, Á., Rodríguez, J.R., Dorostkar, M.M., DeFelipe, J., and Herms, J. (2017). High plasticity of axonal pathology in Alzheimer’s disease mouse models. Acta Neuropathol. Commun. 5, 14.

44. Zhao, S., Zhang, L., Xiang, S., Hu, Y., Wu, Z., and Shen, J. (2022). Gnawing between cells and cells in the immune system: friend or foe? A review of trogocytosis. Front. Immunol. 13, 791006.

45. Tixeira, R., and Poon, I.K.H. (2019). Disassembly of dying cells in diverse organisms. Cell. Mol. Life Sci. 76, 245–257.

46. Vorselen, D. (2022). Dynamics of phagocytosis mediated by phosphatidylserine. Biochem. Soc. Trans. 50, 1281–1291.

47. Sapar, M.L., Ji, H., Wang, B., Poe, A.R., Dubey, K., Ren, X., Ni, J.-Q., and Han, C. (2018). Phosphatidylserine Externalization Results from and Causes Neurite Degeneration in Drosophila. Cell Rep. 24, 2273–2286.

48. von Hofsten, J., and Olsson, P.-E. (2005). Zebrafish sex determination and differentiation: involvement of FTZ-F1 genes. Reprod. Biol. Endocrinol. 3, 63.

49. Peri, F., and Nüsslein-Volhard, C. (2008). Live imaging of neuronal degradation by microglia reveals a role for v0-ATPase a1 in phagosomal fusion in vivo. Cell 133, 916–927.

50. Kim, D.H., Kim, J., Marques, J.C., Grama, A., Hildebrand, D.G.C., Gu, W., Li, J.M., and Robson, D.N. (2017). Pan-neuronal calcium imaging with cellular resolution in freely swimming zebrafish. Nat. Methods 14, 1107–1114.

51. Das, A., and Crump, J.G. (2012). Bmps and id2a act upstream of Twist1 to restrict ectomesenchyme potential of the cranial neural crest. PLoS Genet. 8, e1002710.

52. Barford, K., Keeler, A., McMahon, L., McDaniel, K., Yap, C.C., Deppmann, C.D., and Winckler, B. (2018). Transcytosis of TrkA leads to diversification of dendritic signaling endosomes. Sci. Rep. 8, 4715.

53. Yong, Y., Hughes, C., and Deppmann, C. (2020). A microfluidic culture platform to assess axon degeneration. Methods Mol. Biol. 2143, 83–96.

