## Supplementary materials 1 for "Axonal spheroids are regulated by Schwann cells after peripheral nerve injury"

Spheroid algorithm validation

Sarah Hunter-Chang

2024-10-21

### Purpose

Create a mixed effects model to see if the automated spheroid detection algorithm predicts manual spheroid detection, accounting for repeated measures from the same zebrafish larva at three times: before 1:30, 2:30, and 5:00 post-injury.

### Generating the data set

Automatic Spheroid Detection:

To quantify the number of spheroids over time after injury, time lapse videos of GCaMP6s signal maximum z projections from each injury were cropped so the ROI spanned 150 to 250 µm distal to the injury site. We then used the surface creation module in Bitplane Imaris with a FIJI Labkit machine learning segmentation plugin to detect spheroids using GCaMP6s signal. Specifically, manually identified spheroids were annotated as “foreground” and manually identified non-spheroid signal was annotated as “background.” Multiple iterations of training were performed until additional training stopped improving the algorithm’s abilities to detect spheroids without also detecting false positives. Debris and noise were filtered out of datasets by excluding volumes <1 µm^3 or durations of only 1 frame. The number of spheroids in each larva at each frame was exported from Bitplane - Imaris as a .csv file, then R Studio was used to extract the number of spheroids in each larva from 15 to 300 minutes every 5 minutes post-injury.

Manual Spheroid Detection:

To validate spheroid detection with our Bitplane Imaris FIJI Labkit machine learning segmentation plugin, spheroids from the above videos were manually identified by a blinded reviewer at selected time points: a frame with maximal spheroid signal intensity and morphological variety between 15 and 90 minutes post-injury, at 150 minutes post-injury, and at 300 minutes post-injury.

### Fit the linear mixed-effects model in R Studio

Set up the library

```
library(lme4)
## Loading required package: Matrix
library(tidyverse)
## ── Attaching core tidyverse packages ──────────────────────── tidyverse 2.0.0 ──
## ✔ dplyr     1.1.4     ✔ readr     2.1.5
## ✔ forcats   1.0.0     ✔ stringr   1.5.1
## ✔ ggplot2   3.5.1     ✔ tibble    3.2.1
## ✔ lubridate 1.9.3     ✔ tidyr     1.3.1
## ✔ purrr     1.0.2
## ── Conflicts ────────────────────────────────────────── tidyverse_conflicts() ──
## ✖ tidyr::expand() masks Matrix::expand()
## ✖ dplyr::filter() masks stats::filter()
## ✖ dplyr::lag()    masks stats::lag()
## ✖ tidyr::pack()   masks Matrix::pack()
## ✖ tidyr::unpack() masks Matrix::unpack()
## ℹ Use the conflicted package (<http://conflicted.r-lib.org/>) to force all conflicts to become errors
library(ggeffects)
library(ggplot2)
library(dbscan)
##
## Attaching package: 'dbscan'
##
## The following object is masked from 'package:stats':
##
##     as.dendrogram
library(stargazer)
##
## Please cite as:
##
##  Hlavac, Marek (2022). stargazer: Well-Formatted Regression and Summary Statistics Tables.
##  R package version 5.2.3. https://CRAN.R-project.org/package=stargazer
library(lmerTest)
##
## Attaching package: 'lmerTest'
##
## The following object is masked from 'package:lme4':
##
##     lmer
##
## The following object is masked from 'package:stats':
##
##     step
```

Import the data set

```
my_data <- read_csv("Spheroid_Algorithm_Validation_Raw_Data.csv")
## Rows: 24 Columns: 4
## ── Column specification ────────────────────────────────────────────────────────
## Delimiter: ","
## chr (1): Time
## dbl (3): Subject, Manual_Count, Automated_Count
##
## ℹ Use `spec()` to retrieve the full column specification for this data.
## ℹ Specify the column types or set `show_col_types = FALSE` to quiet this message.
```

Plot the data as an XY plot. X = manual count, Y = automated count. And, for good measure, show which time point the measurement came from by coloring them.

```
my_data %>%
  ggplot(aes(Manual_Count, Automated_Count)) +
  geom_point(aes(color = Time))
```

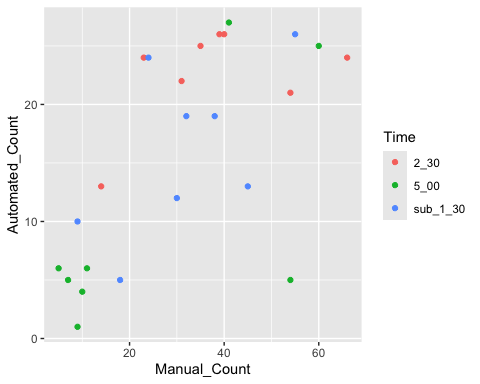

Looks like that one point for the 2:30 group may be an outlier. Let’s try a Local Outlier Factor (LOF) analysis to see:

```
#Get rid of everything but manual and automated count columns in the data frame
outlier_data <- my_data[, 3:4]

# Combine X and Y into a matrix
data_matrix <- as.matrix(outlier_data)

# Apply LOF
lof_scores <- lof(data_matrix, k = 2)  # Adjust k as needed
## Warning in lof(data_matrix, k = 2): lof: k is now deprecated. use minPts = 3
## instead .
# Identify outliers based on LOF scores
outlier_threshold <- 2  # Example threshold
outliers <- outlier_data[lof_scores > outlier_threshold, ]

print(outliers)
## # A tibble: 1 × 2
##   Manual_Count Automated_Count
##          <dbl>           <dbl>
## 1           18               5
```

Visually the identified point does not seem to be an actual outlier, so we will include everything.

Create a mixed effects model with a random effect for each fish to account for effect of each fish

```
model <- lmer(Automated_Count ~ Manual_Count + (1 | Subject), data = my_data)
summary(model)
## Linear mixed model fit by REML. t-tests use Satterthwaite's method [
## lmerModLmerTest]
## Formula: Automated_Count ~ Manual_Count + (1 | Subject)
##    Data: my_data
##
## REML criterion at convergence: 159.6
##
## Scaled residuals:
##      Min       1Q   Median       3Q      Max
## -2.46393 -0.54073 -0.06779  0.60315  1.42408
##
## Random effects:
##  Groups   Name        Variance Std.Dev.
##  Subject  (Intercept) 10.87    3.298
##  Residual             40.16    6.337
## Number of obs: 24, groups:  Subject, 8
##
## Fixed effects:
##              Estimate Std. Error       df t value Pr(>|t|)
## (Intercept)   5.42025    3.22025 11.49646   1.683   0.1193
## Manual_Count  0.34389    0.08668 15.27860   3.967   0.0012 **
## ---
## Signif. codes:  0 '***' 0.001 '**' 0.01 '*' 0.05 '.' 0.1 ' ' 1
##
## Correlation of Fixed Effects:
##             (Intr)
## Manual_Cont -0.841
```

Plot the predicted data from the mixed effects model above as a line against actual data as an xy plot with manual count as x and automated count as y

```
# Plot predicted values using ggeffects as a line, and raw data as scatterplot.
plot(ggpredict(model, terms = c("Manual_Count")), add.data = T)+theme_bw()
## Warning: Argument `add.data` is deprecated and will be removed in the future.
##   Please use `show_data` instead.
## Data points may overlap. Use the `jitter` argument to add some amount of
##   random variation to the location of data points and avoid overplotting.
```

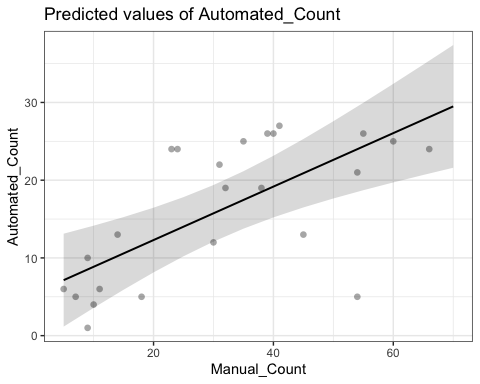

```
ggsave("plot1.tiff")
## Saving 5 x 4 in image
```

Let’s see how good a fit it is:

```
AIC(model)
## [1] 167.6003
```

Let’s check the residuals are normally distributed:

residuals <- residuals(model)
qqnorm(residuals)

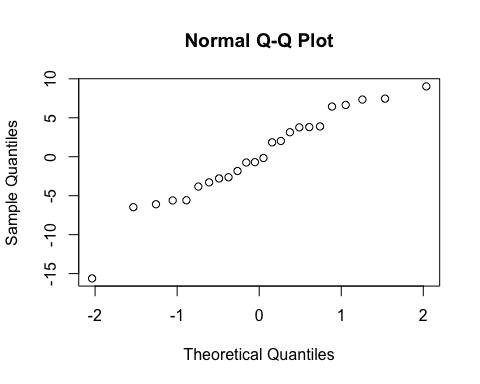

```
shapiro_test <- shapiro.test(residuals)
print(shapiro_test)
##
##  Shapiro-Wilk normality test
##
## data:  residuals
## W = 0.9531, p-value = 0.3158
```

Hooray! They’re normal! The model is good to go.
