## Supplementary materials 2 for "Axonal spheroids are regulated by Schwann cells after peripheral nerve injury"

Spheroid Quantities SC Ablation Poisson

2024-10-21

#### Purpose

Purpose: fit a Poisson mixed effects model to individual zebrafish injury spheroid quantities over time with (Ronidazole, RDZ, treated) and without (Vehicle control, VC, treated) Schwann cell ablation

library(tidyverse)
library(ggeffects)
library(emmeans)
library(lme4)
library(splines)

#### Import the data set

Data should be in long form with a column for time, zebrafish ID, RDZ status, and number of spheroids at each time

mydata <- read_csv("Spheroid_quantities_long.csv")

Remove empty columns

mydata <- mydata[,1:5]

Generate a count of how many observations per fish and per treatment

```
mydata %>%
  count(RDZ, Fish)
## # A tibble: 20 × 3
##    RDZ   Fish      n
##    <chr> <chr> <int>
##  1 RDZ   G        60
##  2 RDZ   H        60
##  3 RDZ   I        60
##  4 RDZ   J        60
##  5 RDZ   K        60
##  6 RDZ   L        60
##  7 RDZ   M        60
##  8 RDZ   N        60
##  9 RDZ   O        60
## 10 RDZ   P        60
## 11 RDZ   Q        60
## 12 RDZ   R        60
## 13 RDZ   S        60
## 14 RDZ   T        60
## 15 VC    A        60
## 16 VC    B        60
## 17 VC    C        60
## 18 VC    D        60
## 19 VC    E        60
## 20 VC    F        60
```

#### Make Time into a minutes variable instead of clock time

First make Time a proper Time variable

```
mydata <- mydata %>%
  drop_na() %>%
  mutate(Time = hms(Time))

mydata
## # A tibble: 1,200 × 5
##    Time     Fish  RDZ   n_spheroids rel_spheroids
##    <Period> <chr> <chr>       <dbl>         <dbl>
##  1 5M 0S    A     VC              0           0
##  2 10M 0S   A     VC              0           0
##  3 15M 0S   A     VC              0           0
##  4 20M 0S   A     VC              0           0
##  5 25M 0S   A     VC              4          11.0
##  6 30M 0S   A     VC              7          19.3
##  7 35M 0S   A     VC             14          38.5
##  8 40M 0S   A     VC             15          41.3
##  9 45M 0S   A     VC             16          44.0
## 10 50M 0S   A     VC             19          52.3
## # ℹ 1,190 more rows
```

Then get the number of minutes past nerve ablation

```
mydata <- mydata %>%
  mutate(Min = as.duration(Time) / dminutes(1))

mydata %>%
  print(n = 30)
## # A tibble: 1,200 × 6
##    Time      Fish  RDZ   n_spheroids rel_spheroids   Min
##    <Period>  <chr> <chr>       <dbl>         <dbl> <dbl>
##  1 5M 0S     A     VC              0           0       5
##  2 10M 0S    A     VC              0           0      10
##  3 15M 0S    A     VC              0           0      15
##  4 20M 0S    A     VC              0           0      20
##  5 25M 0S    A     VC              4          11.0    25
##  6 30M 0S    A     VC              7          19.3    30
##  7 35M 0S    A     VC             14          38.5    35
##  8 40M 0S    A     VC             15          41.3    40
##  9 45M 0S    A     VC             16          44.0    45
## 10 50M 0S    A     VC             19          52.3    50
## 11 55M 0S    A     VC             12          33.0    55
## 12 1H 0M 0S  A     VC             17          46.8    60
## 13 1H 5M 0S  A     VC             23          63.3    65
## 14 1H 10M 0S A     VC             27          74.3    70
## 15 1H 15M 0S A     VC             31          85.3    75
## 16 1H 20M 0S A     VC             28          77.1    80
## 17 1H 25M 0S A     VC             26          71.6    85
## 18 1H 30M 0S A     VC             26          71.6    90
## 19 1H 35M 0S A     VC             26          71.6    95
## 20 1H 40M 0S A     VC             24          66.1   100
## 21 1H 45M 0S A     VC             26          71.6   105
## 22 1H 50M 0S A     VC             23          63.3   110
## 23 1H 55M 0S A     VC             27          74.3   115
## 24 2H 0M 0S  A     VC             33          90.8   120
## 25 2H 5M 0S  A     VC             30          82.6   125
## 26 2H 10M 0S A     VC             35          96.3   130
## 27 2H 15M 0S A     VC             31          85.3   135
## 28 2H 20M 0S A     VC             31          85.3   140
## 29 2H 25M 0S A     VC             29          79.8   145
## 30 2H 30M 0S A     VC             26          71.6   150
## # ℹ 1,170 more rows
```

Yes, this worked

Update 2024-10-18. SHC says “data set should start at 15 minutes, not 5. I was looking over things again and remembered that some of my time lapses started >10 minutes after injury. I’d automatically filled every empty time point with a 0 because the spheroid detection software just leaves time points in which it detects 0 spheroids blank, even if the time lapse included that time point. So now some of the spheroid quantities from <15 minutes were actually unknown, but erroneously entered as 0”

Subset for 15+ minutes

```
mydata <- mydata %>%
  filter(Min >= 15)
```

Now regenerate the count of how many observations per fish and per treatment

```
mydata %>%
  count(RDZ, Fish)
## # A tibble: 20 × 3
##    RDZ   Fish      n
##    <chr> <chr> <int>
##  1 RDZ   G        58
##  2 RDZ   H        58
##  3 RDZ   I        58
##  4 RDZ   J        58
##  5 RDZ   K        58
##  6 RDZ   L        58
##  7 RDZ   M        58
##  8 RDZ   N        58
##  9 RDZ   O        58
## 10 RDZ   P        58
## 11 RDZ   Q        58
## 12 RDZ   R        58
## 13 RDZ   S        58
## 14 RDZ   T        58
## 15 VC    A        58
## 16 VC    B        58
## 17 VC    C        58
## 18 VC    D        58
## 19 VC    E        58
## 20 VC    F        58
```

#### Plots

Make a spaghetti plot showing **number of spheroids** over time for each individual fish

```
mydata %>%
  ggplot(aes(Min, n_spheroids)) +
  geom_line(aes(group = Fish)) +
  facet_wrap(vars(RDZ))
```

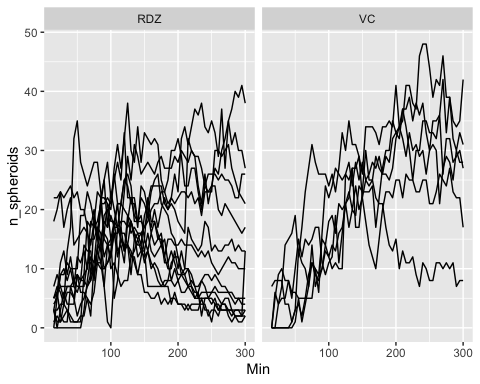

```
# see one fish
mydata %>%
  filter(Fish == "A") %>%
  ggplot(aes(Min, n_spheroids)) +
  geom_line(aes(group = Fish)) +
  facet_wrap(vars(RDZ))
```

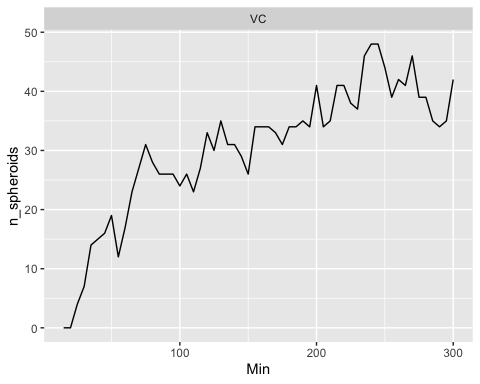

Regenerate the plot from Prism

Calculate the average n_spheroid at each time point for each drug treatment group. Put VC first.

```
mydataSUM <- mydata %>%
  group_by(RDZ, Min) %>%
  summarize(mean_spheroids = mean(n_spheroids),
            sem_spheroids = sd(n_spheroids) / sqrt(n()),
            lower = mean_spheroids - sem_spheroids,
            upper = mean_spheroids + sem_spheroids,
            .groups = "drop") %>%
  mutate(RDZ = factor(RDZ, levels = c("VC", "RDZ")))

mydataSUM
## # A tibble: 116 × 6
##    RDZ     Min mean_spheroids sem_spheroids lower upper
##    <fct> <dbl>          <dbl>         <dbl> <dbl> <dbl>
##  1 RDZ      15           5.64          1.72  3.92  7.36
##  2 RDZ      20           6.57          1.79  4.78  8.36
##  3 RDZ      25           7.57          1.97  5.61  9.54
##  4 RDZ      30           7.43          1.68  5.75  9.11
##  5 RDZ      35           8.21          1.80  6.42 10.0
##  6 RDZ      40           9.21          1.97  7.24 11.2
##  7 RDZ      45           8.86          2.34  6.51 11.2
##  8 RDZ      50           9.93          2.54  7.39 12.5
##  9 RDZ      55           9.86          2.17  7.68 12.0
## 10 RDZ      60          11.3           1.90  9.39 13.2
## # ℹ 106 more rows
```

Now plot

```
mydataSUM %>%
  ggplot(aes(Min, mean_spheroids, color = RDZ, fill = RDZ)) +
  geom_line(linewidth = 2) +
  geom_ribbon(aes(ymax = upper, ymin = lower),
              alpha = .2,
              color = NA)
```

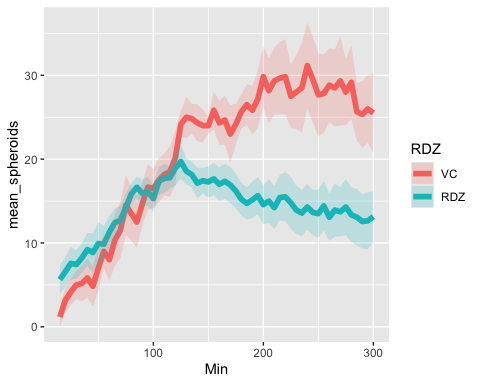

#### Fit Poisson mixed effects model

```
mod1 <- glmer(n_spheroids ~ Min * RDZ + (1|Fish), family = poisson, data = mydata)
summary(mod1, corr = FALSE)
## Generalized linear mixed model fit by maximum likelihood (Laplace
##   Approximation) [glmerMod]
##  Family: poisson  ( log )
## Formula: n_spheroids ~ Min * RDZ + (1 | Fish)
##    Data: mydata
##
##      AIC      BIC   logLik deviance df.resid
##   9207.0   9232.3  -4598.5   9197.0     1155
##
## Scaled residuals:
##     Min      1Q  Median      3Q     Max
## -4.8321 -1.2831  0.0761  1.1734  7.8214
##
## Random effects:
##  Groups Name        Variance Std.Dev.
##  Fish   (Intercept) 0.1387   0.3725
## Number of obs: 1160, groups:  Fish, 20
##
## Fixed effects:
##               Estimate Std. Error z value Pr(>|z|)
## (Intercept)  2.4341882  0.1017092  23.933  < 2e-16 ***
## Min          0.0008564  0.0001115   7.682 1.57e-14 ***
## RDZVC       -0.2420752  0.1854430  -1.305    0.192
## Min:RDZVC    0.0037607  0.0001843  20.405  < 2e-16 ***
## ---
## Signif. codes:  0 '***' 0.001 '**' 0.01 '*' 0.05 '.' 0.1 ' ' 1
## optimizer (Nelder_Mead) convergence code: 0 (OK)
## Model is nearly unidentifiable: very large eigenvalue
##  - Rescale variables?
## Model is nearly unidentifiable: large eigenvalue ratio
##  - Rescale variables?
```

See the model in a plot

plot(ggpredict(mod1, terms = c("Min", "RDZ")))

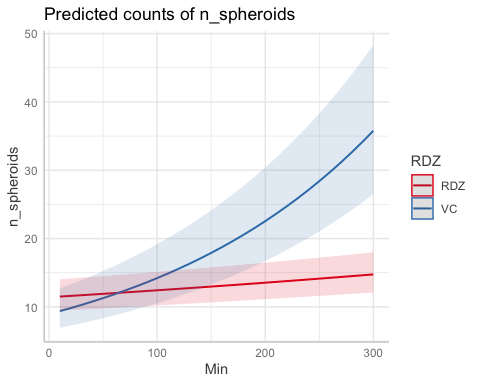

```
# add the raw data
plot(ggpredict(mod1, terms = c("Min", "RDZ")),
     add.data = T)
```

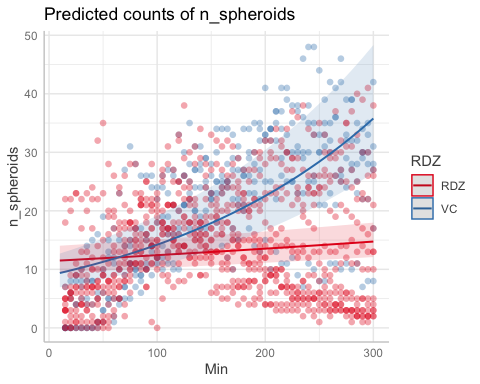

The RDZ do not seem to fit well

See the model for each fish

```
ggpredict(mod1, terms = c("Min", "RDZ", "Fish"), type = "random") %>%
  as_tibble() %>%
  rename(Min = x,
         RDZ = group,
         Fish = facet) %>%
  ggplot() +
  geom_line(aes(x = Min, y = predicted, color = RDZ)) +
  facet_wrap(~Fish)
```

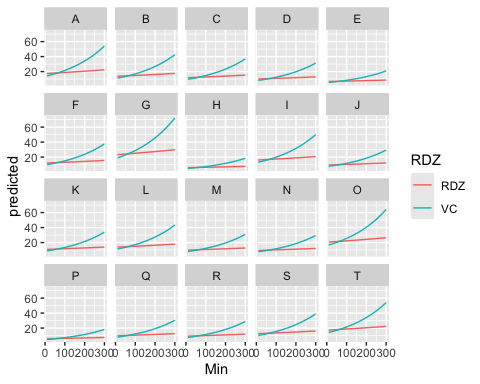

```
# add the raw data to see the fit
ggpredict(mod1, terms = c("Min", "RDZ", "Fish"), type = "random") %>%
  as_tibble() %>%
  rename(Min = x,
         RDZ = group,
         Fish = facet) %>%
  ggplot() +
  geom_line(aes(x = Min, y = predicted, color = RDZ)) +
  facet_wrap(~Fish) +
  geom_line(data = mydata, aes(Min, n_spheroids, group = Fish, color = RDZ))
```

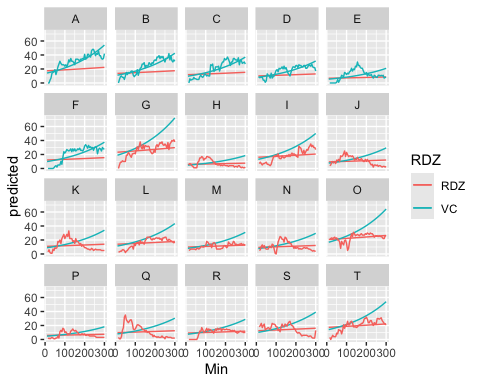

Actually, this model fits better than I thought.

### Poisson mixed effects model with restricted cubic splines

Try with non-linear spline terms in case that complexity is better

```
mod2 <- glmer(n_spheroids ~ ns(Min, df = 3) + RDZ + (1|Fish),
              family = poisson,
              data = mydata)
```

See the model

plot(ggpredict(mod2, terms = c("Min [all]", "RDZ")))

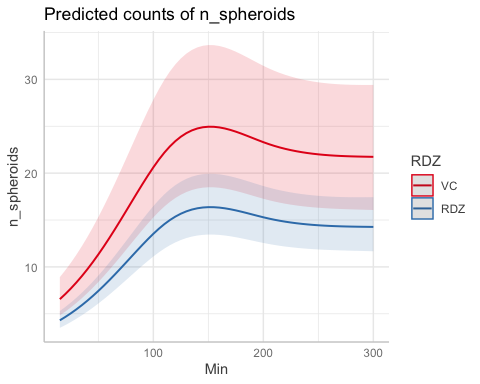

See this model for each fish

```
ggpredict(mod2, terms = c("Min [all]", "RDZ", "Fish"), type = "random") %>%
  as_tibble() %>%
  rename(Min = x,
         RDZ = group,
         Fish = facet) %>%
  ggplot() +
  geom_line(aes(x = Min, y = predicted, color = RDZ)) +
  facet_wrap(~Fish)
```

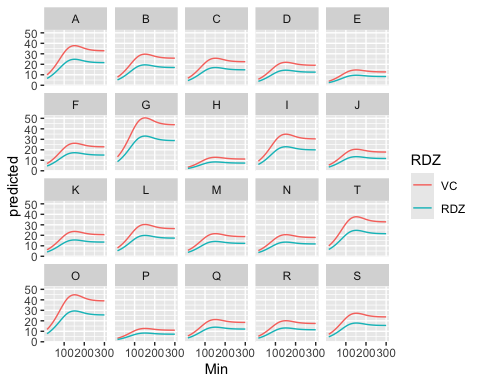

```
# add the raw data to see the fit
ggpredict(mod2, terms = c("Min [all]", "RDZ", "Fish"), type = "random") %>%
  as_tibble() %>%
  rename(Min = x,
         RDZ = group,
         Fish = facet) %>%
  ggplot() +
  geom_line(aes(x = Min, y = predicted, color = RDZ)) +
  facet_wrap(~Fish) +
  geom_line(data = mydata, aes(Min, n_spheroids, group = Fish, color = RDZ))
```

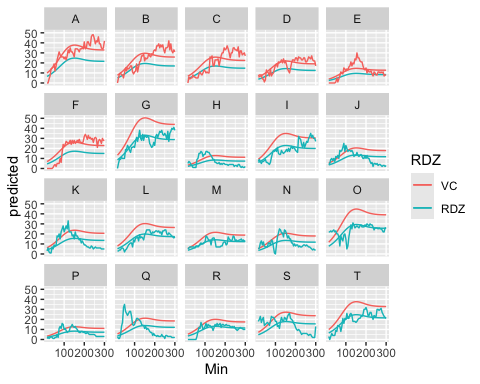

Hmm, not sure this version is better

```
AIC(mod1, mod2)
##      df      AIC
## mod1  5 9207.010
## mod2  6 8719.908
```

According to AIC, mod2 does have a better fit

#### Try splines with an interaction term

```
mod3 <- glmer(n_spheroids ~ ns(Min, df = 3) * RDZ + (1|Fish),
              family = poisson,
              data = mydata)
```

See the model

plot(ggpredict(mod3, terms = c("Min [all]", "RDZ")))

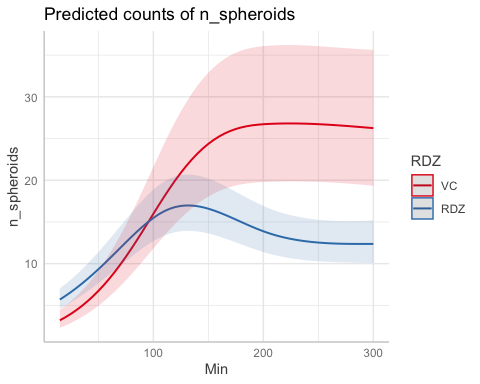

Ahh, this might be better

See this model for each fish

```
ggpredict(mod3, terms = c("Min [all]", "RDZ", "Fish"), type = "random") %>%
  as_tibble() %>%
  rename(Min = x,
         RDZ = group,
         Fish = facet) %>%
  ggplot() +
  geom_line(aes(x = Min, y = predicted, color = RDZ)) +
  facet_wrap(~Fish)
```

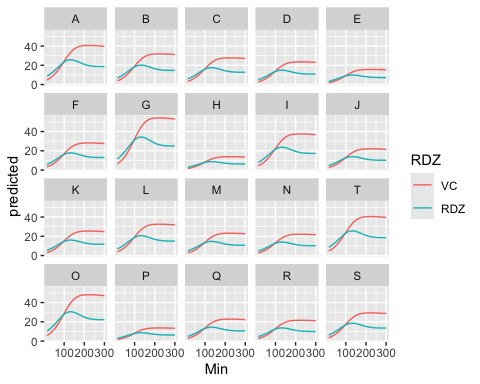

```
# add the raw data to see the fit
ggpredict(mod3, terms = c("Min [all]", "RDZ", "Fish"), type = "random") %>%
  as_tibble() %>%
  rename(Min = x,
         RDZ = group,
         Fish = facet) %>%
  ggplot() +
  geom_line(aes(x = Min, y = predicted, color = RDZ)) +
  facet_wrap(~Fish) +
  geom_line(data = mydata, aes(Min, n_spheroids, group = Fish, color = RDZ))
```

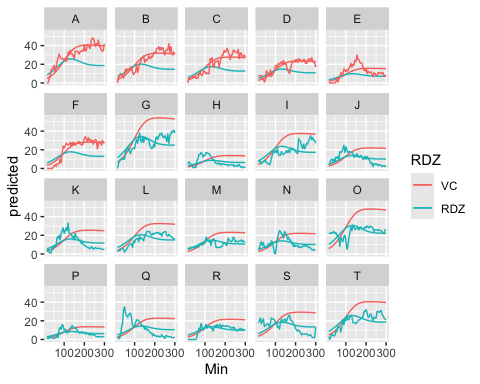

Check out the difference in AIC between these mod2 and mod3

```
AIC(mod2, mod3)
##      df      AIC
## mod2  6 8719.908
## mod3  9 8157.726
```

Mod3 is preferred

See the average raw data plotted with this model

```
ggpredict(mod3, terms = c("Min [all]", "RDZ")) %>%
  as_tibble() %>%
  rename(Min = x,
         RDZ = group) %>%
  ggplot() +
  geom_line(aes(x = Min, y = predicted, color = RDZ),
            linewidth = 2) +
  geom_line(data = mydataSUM, aes(Min, mean_spheroids, color = RDZ)) +
  geom_ribbon(data = mydataSUM, aes(Min, mean_spheroids, fill = RDZ,
                                    ymax = upper, ymin = lower),
              alpha = .2,
              color = NA) +  # Ribbon with fill by RDZ
  scale_color_manual(values = c("RDZ" = "#FFCC66", "VC" = "#33CCFF")) +  # Assign specific colors to RDZ
  scale_fill_manual(values = c("RDZ" = "#FFCC66", "VC" = "#33CCFF")) +  # Same colors for ribbon fill
theme_classic() +
  theme(axis.line = element_line(size = 1))
```

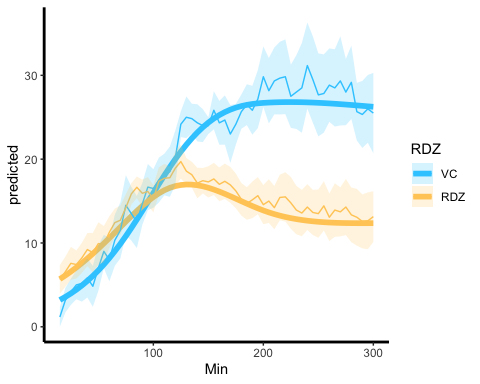

ggsave("plot1.tiff", width = 8, height = 6, dpi = 300)

This model fits these data pretty FREAKIN WELL! I am JAZZED!

See the summary

```
summary(mod3, corr = FALSE)
## Generalized linear mixed model fit by maximum likelihood (Laplace
##   Approximation) [glmerMod]
##  Family: poisson  ( log )
## Formula: n_spheroids ~ ns(Min, df = 3) * RDZ + (1 | Fish)
##    Data: mydata
##
##      AIC      BIC   logLik deviance df.resid
##   8157.7   8203.2  -4069.9   8139.7     1151
##
## Scaled residuals:
##     Min      1Q  Median      3Q     Max
## -4.4111 -1.1783 -0.0869  0.9143  9.6180
##
## Random effects:
##  Groups Name        Variance Std.Dev.
##  Fish   (Intercept) 0.1388   0.3725
## Number of obs: 1160, groups:  Fish, 20
##
## Fixed effects:
##                         Estimate Std. Error z value Pr(>|z|)
## (Intercept)             1.739196   0.107505  16.178   <2e-16 ***
## ns(Min, df = 3)1        0.381829   0.039904   9.569   <2e-16 ***
## ns(Min, df = 3)2        1.944269   0.095240  20.414   <2e-16 ***
## ns(Min, df = 3)3       -0.004215   0.033125  -0.127   0.8988
## RDZVC                  -0.575611   0.200510  -2.871   0.0041 **
## ns(Min, df = 3)1:RDZVC  1.181737   0.069013  17.123   <2e-16 ***
## ns(Min, df = 3)2:RDZVC  1.913358   0.190310  10.054   <2e-16 ***
## ns(Min, df = 3)3:RDZVC  0.947985   0.052985  17.891   <2e-16 ***
## ---
## Signif. codes:  0 '***' 0.001 '**' 0.01 '*' 0.05 '.' 0.1 ' ' 1
```

Ok, pretty much everything is significant. See the predicted average differences between RDZ and VC at given times

every hour?

```
60*1:5
## [1]  60 120 180 240 300
```

See the predicted number of spheroids at given time points

```
emm <- emmeans(mod3, ~RDZ | Min,
        at = list(Min =  c(60, 120, 180, 240, 300)),
        type = "response")

emm
## Min =  60:
##  RDZ rate   SE  df asymp.LCL asymp.UCL
##  RDZ 10.6 1.07 Inf      8.68      12.9
##  VC   8.2 1.28 Inf      6.04      11.1
##
## Min = 120:
##  RDZ rate   SE  df asymp.LCL asymp.UCL
##  RDZ 16.8 1.69 Inf     13.77      20.5
##  VC  20.0 3.08 Inf     14.79      27.0
##
## Min = 180:
##  RDZ rate   SE  df asymp.LCL asymp.UCL
##  RDZ 15.0 1.51 Inf     12.30      18.3
##  VC  26.3 4.03 Inf     19.50      35.5
##
## Min = 240:
##  RDZ rate   SE  df asymp.LCL asymp.UCL
##  RDZ 12.7 1.28 Inf     10.39      15.4
##  VC  26.8 4.10 Inf     19.83      36.1
##
## Min = 300:
##  RDZ rate   SE  df asymp.LCL asymp.UCL
##  RDZ 12.4 1.29 Inf     10.07      15.2
##  VC  26.2 4.10 Inf     19.33      35.6
##
## Confidence level used: 0.95
## Intervals are back-transformed from the log scale
```

Are the drug treatment groups different at each timepoint?

```
pairs(emm)
## Min = 60:
##  contrast ratio     SE  df null z.ratio p.value
##  RDZ / VC 1.291 0.2397 Inf    1   1.376  0.1689
##
## Min = 120:
##  contrast ratio     SE  df null z.ratio p.value
##  RDZ / VC 0.839 0.1545 Inf    1  -0.952  0.3411
##
## Min = 180:
##  contrast ratio     SE  df null z.ratio p.value
##  RDZ / VC 0.569 0.1043 Inf    1  -3.074  0.0021
##
## Min = 240:
##  contrast ratio     SE  df null z.ratio p.value
##  RDZ / VC 0.473 0.0867 Inf    1  -4.086  <.0001
##
## Min = 300:
##  contrast ratio     SE  df null z.ratio p.value
##  RDZ / VC 0.471 0.0885 Inf    1  -4.007  0.0001
##
## Tests are performed on the log scale
```
