## Supplementary figure 1 for "Axonal spheroids are regulated by Schwann cells after peripheral nerve injury"

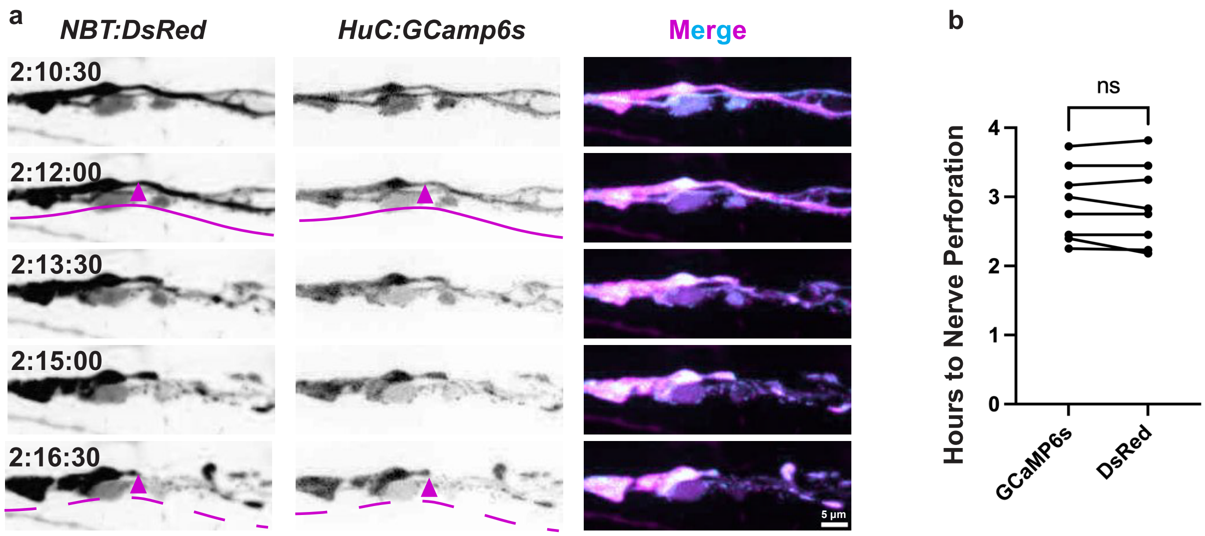


**Supplementary Figure 1:** GCaMP6s is a valid marker of nerve perforation. a) Intact (solid lines) and perforated (dashed lines) axons (arrowheads) detected by *NBT:DsRed* (left column) and by *HuC:GCaMP6s* (middle column) do not significantly differ in time to nerve perforation (b, N=7 larvae, p=0.6250, two-tailed Wilcoxon matched-pairs signed rank test).
