## Supplementary figure 2 for "Axonal spheroids are regulated by Schwann cells after peripheral nerve injury"

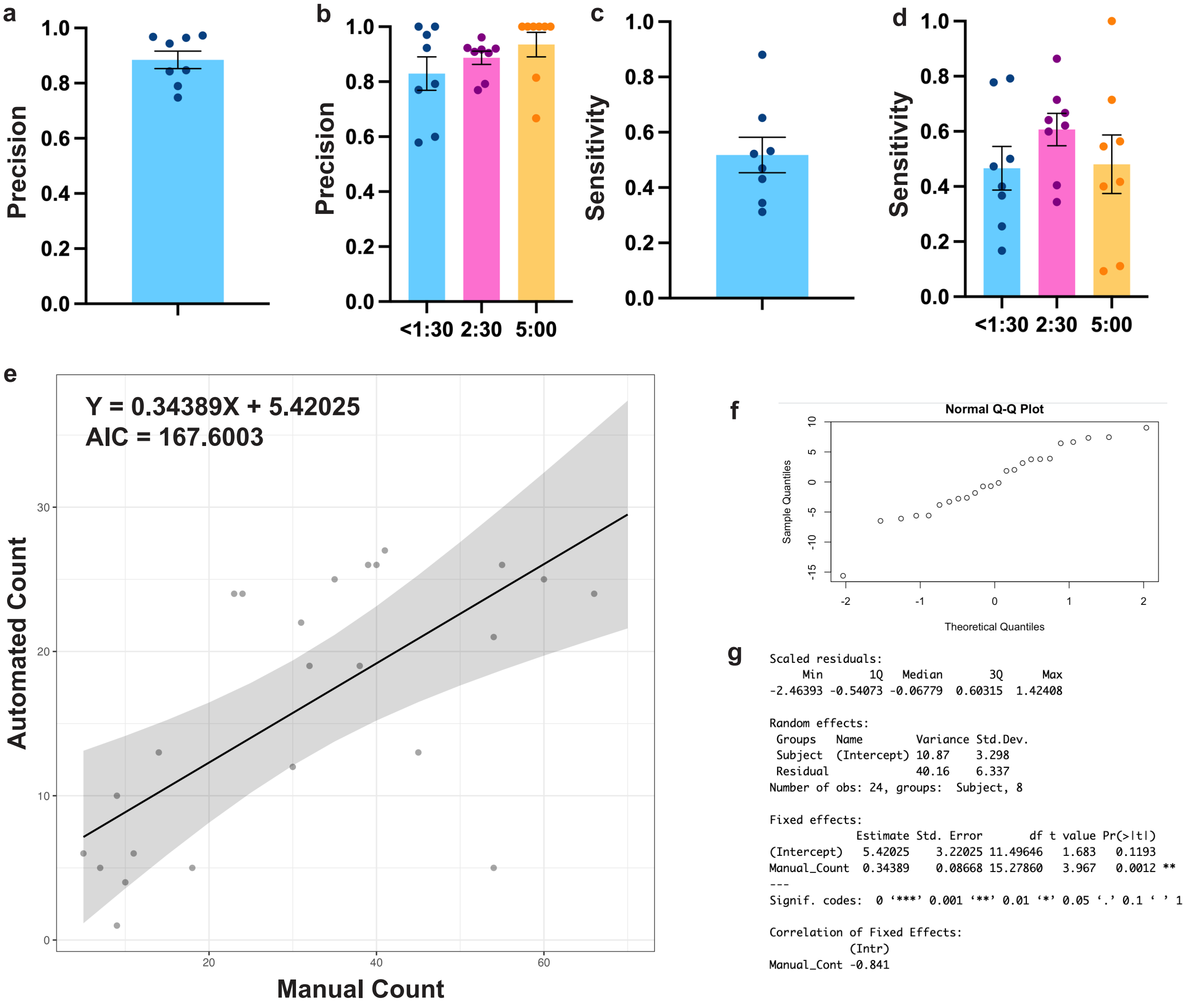


**Supplementary Figure 2: Automated spheroid detection predicts manually detected spheroid quantity.** a) Automated spheroid detection precision averaged from each larva (N=8 larvae). A blinded reviewer manually identified spheroids in algorithm training videos at selected time points: a frame with maximal spheroid signal intensity and morphological variety between 15 and 90 minutes post-injury, at 150 minutes post-injury, and at 300 minutes post-injury. Spheroids detected by the algorithm were considered true positives (*TP*), those detected only by the automated method were false positives (*FP*), and those detected only by manual method were false negatives (*FN*). Precision was calculated as *TP* / (*TP* + *FP*). b) Precision at each time after injury (N=8 larvae, p=0.2354, Friedman test). c) Automated spheroid detection sensitivity from each larva (N=8 larvae). Sensitivity was calculated as *TP* / (*TP* + *FN*). d) Sensitivity at each time after injury (N=8 larvae, p=0.2823, one-way ANOVA). e) Manual versus automated spheroid counts at <1:30, 2:30, and 5:00 hours post-injury (points) fit with a linear mixed effect model to take into account repeated measurements from each fish (line, Supplementary Materials 1). f) A normal Q-Q plot showing residuals from the observed values to the predicted values in e. g) A summary of the linear mixed effect model in g. For every manually detected spheroid, 0.34389 spheroids were automatically detected (N=8 larvae, 3 time points from each larva, p=0.0012, Satterthwaite’s method)
