## Supplementary figure 3 for "Axonal spheroids are regulated by Schwann cells after peripheral nerve injury"

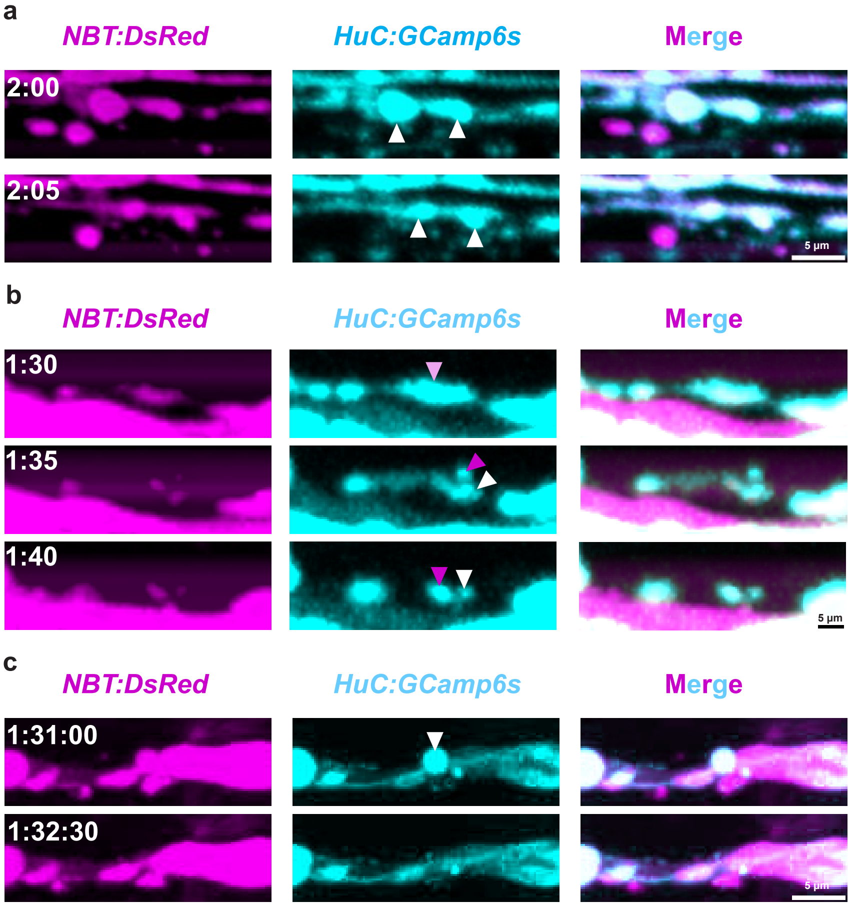


**Supplementary Figure 3: Spheroid fates are visible with both *HuC:GCaMP6s* and *NBT:DsRed* labeling.** a) Spheroids shrinking (white arrowsheads) in both the neuronal calcium (*HuC:GCaMP6s*) and cytosol (*NBT:DsRed*) channels (N=6 larvae). b) A spheroid (pink arrowhead) fragments (magenta and white arrowheads) visibly with both neuronal calcium (*HuC:GCaMP6s*) and cytosol (*NBT:DsRed*) labeling (N=6 larvae). c) Uniform spheroid disappearance (white arrowhead) observed in both the neuronal calcium (*HuC:GCaMP6s*) and cytosol (*NBT:DsRed*) channels (N=6 larvae).
