## Supplementary material for "Axonal spheroids are regulated by Schwann cells after peripheral nerve injury": Captions

**Supplementary video 1: Spheroids shrink.** A *HuC:GCaMP6s*-labeled spheroid (arrowhead) shrinks after pLLN axotomy.

**Supplementary video 2: Spheroids break down.** A *HuC:GCaMP6s*-labeled spheroid (arrowhead) undergoes a series of morphological changes, breaking down from one uniform spherical shape into multiple smaller bodies.

**Supplementary video 3: Spheroids undergo uniform disappearance.** A *HuC:GCaMP6s*-labeled spheroid (arrowhead) disappears uniformly, or without any apparent morphological change.

**Supplementary video 4: Schwann cell interactions with spheroids occur intracellularly.** *HuC:GCamp6s*- (cyan) labeled spheroids (arrowheads) move off of their axons and inside of Schwann cell cytoplasm (*sox10:Gal4;UAS:NTR-mCherry,* yellow).

**Supplementary video 5: Spheroids expose phosphatidylserine *in vivo.*** Phosphatidylserine-labeled spheroids appear in a beads-on-a-string pattern (arrowheads in phosphatidylserine channel). *HuC:jRGECO1b* (cyan)- labeled spheroids are surrounded by phosphatidylserine (*Et(s1101:Gal4);UAS:SecA5-YFP*, yellow, arrowheads in merged channel).

**Supplementary video 6: Spheroids persist in the presence of Schwann cells.** *HuC:GCaMP6s*-labeled spheroids persist as smaller, broken down spheroids out to later times after pLLN injury in a vehicle control-treated (*sox10:Gal4;UAS:NTR*-mCherry) larva.

**Supplementary video 7: Spheroids do not persist as long and undergo uniform disappearance more in the absence of Schwann cells.** *HuC:GCaMP6s*-labeled spheroids disappear more uniformly and more rapidly after pLLN injury in a Ronidazole-treated (*sox10:Gal4;UAS:NTR*-mCherry) larva.
